# Genetic Architecture of Complex Traits and Disease Risk Predictors

**DOI:** 10.1101/2020.02.12.946608

**Authors:** Soke Yuen Yong, Timothy G. Raben, Louis Lello, Stephen D.H. Hsu

## Abstract

Genomic prediction of complex human traits (e.g., height, cognitive ability, bone density) and disease risks (e.g., breast cancer, diabetes, heart disease, atrial fibrillation) has advanced considerably in recent years. Predictors have been constructed using penalized algorithms that favor sparsity: i.e., which use as few genetic variants as possible. We analyze the specific genetic variants (SNPs) utilized in these predictors, which can vary from dozens to as many as thirty thousand. We find that the fraction of SNPs in or near genic regions varies widely by phenotype. For the majority of disease conditions studied, *a large amount* of the variance is accounted for by SNPs outside of coding regions. The state of these SNPs cannot be determined from exome-sequencing data. This suggests that exome data alone will miss much of the heritability for these traits – i.e., existing PRS cannot be computed from exome data alone. We also study the fraction of SNPs and of variance that is in common between pairs of predictors. The DNA regions used in disease risk predictors so far constructed seem to be largely disjoint (with a few interesting exceptions), suggesting that individual genetic disease risks are largely uncorrelated. It seems possible in theory for an individual to be a low-risk outlier in all conditions simultaneously.

## Introduction

Genomic prediction of complex traits and disease risks has advanced considerably thanks to the recent advent of large data sets and improved algorithms. These algorithms range from simple regression, applied to one SNP at a time to estimate statistical significance and effect size (e.g., as used in GWAS), to high dimensional optimization methods such as compressed sensing or sparse learning [1–4]. They produce Polygenic Risk Scores (PRS) or Polygenic Scores (PGS): functions that map the state of an individual’s DNA at specific locations (SNPs), to a risk score or predicted quantitative trait value.

Predictors (PGS or PRS) now exist for a number of important traits and risks, many of which have undergone out of sample testing (i.e., validation in groups of individuals not used in training and from other data sets or from separate ancestries.)[5–7]. The genetic architectures uncovered vary significantly: the number of SNPs required to capture most of the predictor variance ranges from a few dozen to many thousands. In contrast, traditional Genome Wide Association studies (GWAS) can implicate the entire genome [8, 9], making them unwieldy to analyze.

In the case of disease risk, the predictors are already good enough to identify *risk outliers*. That is, individuals with unusually high (or low) risk of a specific condition. There are many clinical applications for such predictors [5, 10–19] (although there is still much work to be done to overcome sampling and algorithmic biases and disparity [20, 21]). Below we mention two examples.

### Breast Cancer

Certain variants in the BRCA1 and BRCA2 genes are known to elevate Breast Cancer risk significantly [22, 23]. However, these mutations affect no more than a few women per thousand in the general population[24–26]. By contrast, PRS using thousands of common SNP variants can now identify an order of *ten times* as many women who are in the high-risk category [5, 7, 10, 27, 28]. Standard of Care for high-risk women typically includes additional screening, such as mammograms beginning a decade earlier than for normal risk women. Early detection can also lead to significant cost savings [29]. What can we say about the thousands of common SNPs used in the PRS? Do they overlap with SNPs used in PRS for other conditions (e.g., other cancers)?

### Height

Idiopathic Short Stature (ISS) refers to extreme short stature that does not have a diagnostic explanation (e.g., height below 5 foot 2 inches in adult males)[30]. Growth hormone treatment is sometimes prescribed for children who are at risk for this condition, at a cost in the $100k range. Typically, these would be children in the bottom percentiles for height within their age group [31, 32]. However, it is difficult for pediatric endocrinologists, whose responsibility it is to prescribe HGH for these children, to know whether the child is simply passing through a temporary phase of slow growth (and will, by adulthood, reach normal height)[33, 34]. Adult height prediction from DNA (with 95 percent confidence interval roughly ±2 inches) will allow physicians to avoid expensive HGH treatment (with significant potential side-effects) for children who are merely short for their age (late-developing) and are likely to be in the normal range in adulthood.

For the first time, we can begin to address some general questions concerning the genetic architectures of complex traits. In this paper we address the following questions:

1. What is the (qualitative) genetic architecture of specific disease risks? How many SNPs, where are they, how many genes?
2. How much of the total risk is controlled by loci in coding vs non-coding regions?
3. Is exome sequencing data sufficient for computation of Polygenic Risk Scores (PRS)?
4. How much genetic overlap exists between different disease architectures? With millions of SNPs in the genome it is entirely possible that different diseases have nearly disjoint genetic architectures – i.e., risk is mostly controlled by distinct regions of DNA. On the other hand, we might uncover overlap regions of DNA which affect multiple disease risks simultaneously.

In this paper we consider predictors for a selection of disease conditions/traits: asthma, atrial fibrillation, basal cell carcinoma, breast cancer, coronary artery disease, type-1 diabetes, type-2 diabetes, diastolic blood pressure, educational years, gallstones, glaucoma, gout, heart attack, height, high cholesterol, hypertension, hypothyroidism, malignant melanoma, menopause, pulse rate, and systolic blood pressure. All predictors, except the coronary artery disease predictor, were built by training on case-control phenotype data from the UK Biobank [35, 36] that relied on custom array genotyping (see Appendix C for details). This array was designed to have detailed coverage in areas known to be associated with certain phenotypes, and to contain a wide sampling of the entire human genome. More details of the array design can be found on the UK Biobank website https://www.ukbiobank.ac.uk/scientists-3/uk-biobank-axiom-array/ and in Appendix C. These predictors were derived using the L1 penalized regression (sparse learning) methods found in [5, 6]. Recent algorithmic benchmarking for complex trait prediction in plants and animals has shown that linear methods work as well if not better than non-linear, Bayesian, or deep learning approaches for current data set sizes [37]. Predictors for diastolic blood pressure, systolic blood pressure, and pulse rate are reported for the first time here, but were designed as described in [5]. The coronary artery predictor originated from [7], and the associated minor allele frequencies were obtained using Ensembl’s minor allele frequency calculator [38].

Of course, not all of the genetic variants affecting disease risk have been discovered. The PRS continue to improve as more training data become available. However, the SNPs used in the existing PRS tend to be those that account for the most risk variance. Equivalently, the statistical evidence supporting their association with the disease risk is highest. Relevant SNPs that have yet to be discovered are either common SNPs with very small effect size, or very rare SNPs that are not probed using existing gene arrays.

## Variance and Effect Sizes

The majority of this work deals with characterizing the relative sizes of the effect of particular SNPs on the performance of a polygenic predictor. A predictor in this case is a set of weights, {*β*_*i*_}, for a set of SNPs, *S*. An individual’s phenotype, *y*, can be described by

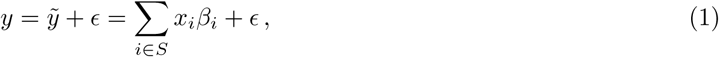

where *x*_*i*_∈{0, 1, 2} is the number of minor alleles for SNP *i*,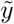 is the predicted value of the phenotype, and *ϵ* is an error term. Our primary object of interest is the variance of this prediction. The contribution of a single SNP to this variance is expressed in terms of the *β*_*i*_ and the minor allele frequency (MAF), *f*_*i*_, as:

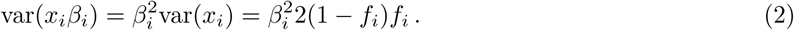

In the limit of small MAF, this becomes 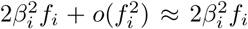. The overall variance of the individual’s predicted phenotype can thus be described as:

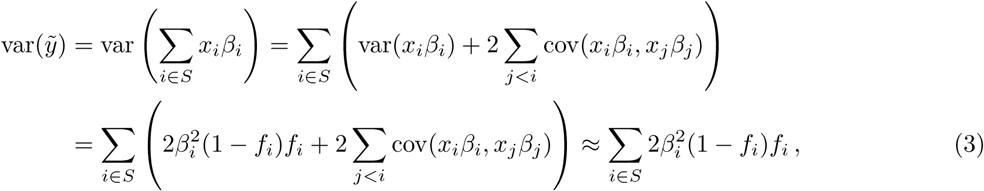

where the final approximation holds when the SNPs are largely uncorrelated. This is true for most minor allele frequencies, and has been checked empirically in particular instances (see for example [6]). For the predictors in this work, there are only a handful of SNPs with correlation of about 0.01 lying within 2000 kilo base pairs of each other. In this sense, the variance due to each SNP can be considered as a linear effect. We can then calculate the *variance accounted for* by a subset, *𝒥* ⊂ *S*, of the predictor SNPs, as a percentage of the total variance of the individual’s predicted phenotype:

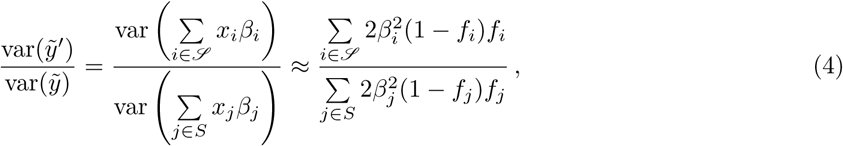

where again, the final approximation holds when the SNPs are uncorrelated.

## Results and Analysis

### Predictor SNPs in genic regions

We are interested in investigating how predictor SNPs located in genic regions impact our predictors.

The most obvious way to identify predictor SNPs located inside genic regions is to define a SNP as being within a genic region if its genomic coordinates fall between the start-point and end-point coordinates of any protein-coding gene. The GENCODE Release 19 annotation of the human genome [39] (based on the GRCh37.p13 reference human genome assembly [40]) was chosen as the source of our reference set of gene boundary coordinates. Currently, it is still unclear where exactly genic regions end and intergenic regions begin [41–43], and so there is a possibility that this choice of gene boundary coordinates may not be definitive for our purposes. We asked these questions: as these reference gene boundary coordinates are varied, by how much does the separation of predictor SNPs into genic and non-genic categories vary, and how significantly does the influence of the (increasingly large) genic section of the predictor SNPs change?

Figure 1 shows for a selection of disease conditions how the number of the predictor SNPs categorized as located in genic regions - expressed as a percentage of the total number of predictor SNPs for that disease condition - changes as all gene boundaries (according to GENCODE Release 19) are expanded by an increasing number *k* of kilo base pairs at both ends. At the reference gene boundaries (*k* = 0), the percentage of predictor SNPs which are genic ranges from about 50% (many disease conditions) to about 60% (gallstones, malignant melanoma, atrial fibrillation); while at *k* = 30, this percentage rises to between 60% (breast cancer, type-1 diabetes, education years) to 75% (gallstones, malignant melanoma). This description excludes the coronary artery disease predictor (42.5% to 55%), which has a distinctly low proportion of genic predictor SNPs. For all disease conditions, the increase in the genic percentage of predictor SNPs with *k* occurs at roughly the same rate, which appears to be almost linear. This suggests that the predictor SNPs for every disease condition are approximately uniformly distributed by distance outside the reference gene boundary coordinates.

**Figure 1:**
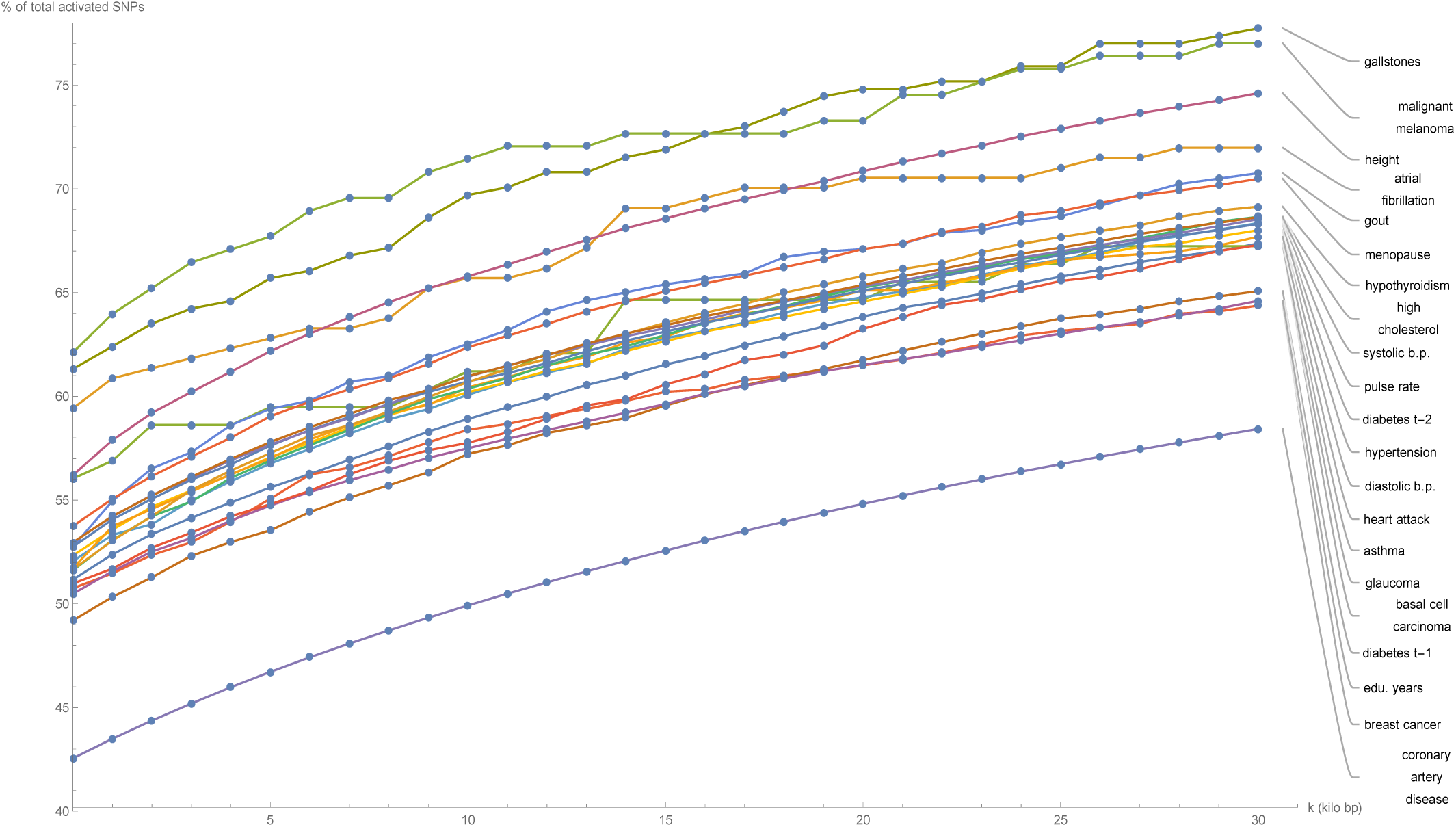
Plots of the number of predictor SNPs located within genic regions, expressed as a percentage of the total number of predictor SNPs for that disease condition, against expansion of GENCODE Release 19 gene boundaries by k kilo base pairs.

Figure 2 shows for each disease condition how the variance accounted for by the predictor SNPs located in genic regions - expressed as a percentage of the total variance accounted for by all predictor SNPs for that condition - changes as the boundaries of every gene (according to GENCODE Release 19) are expanded by *k* kilo base pairs at both ends. At the reference gene boundaries (*k* = 0), the percentage of predictor variance accounted for by SNPs in genic regions ranges from about 40% (breast cancer, type-1 diabetes) to 90% (gallstones, gout, malignant melanoma), with notable outliers at 25% (atrial fibrillation) and 20% (coronary artery disease). For the majority of disease conditions, the percentage of variance accounted for by the genic predictor SNPs remains approximately flat as *k* is increased, meaning that practically all the variance accounted for by genic predictor SNPs is due to SNPs contained within the reference gene boundaries. Noticeable exceptions occur for glaucoma at *k* = 6.5, breast cancer at *k* = 17.5, and menopause at *k* = 27.5, where the observed large jumps in variance indicate the presence of some SNP(s) at that genomic location with a significant effect on that specific disease condition.

**Figure 2:**
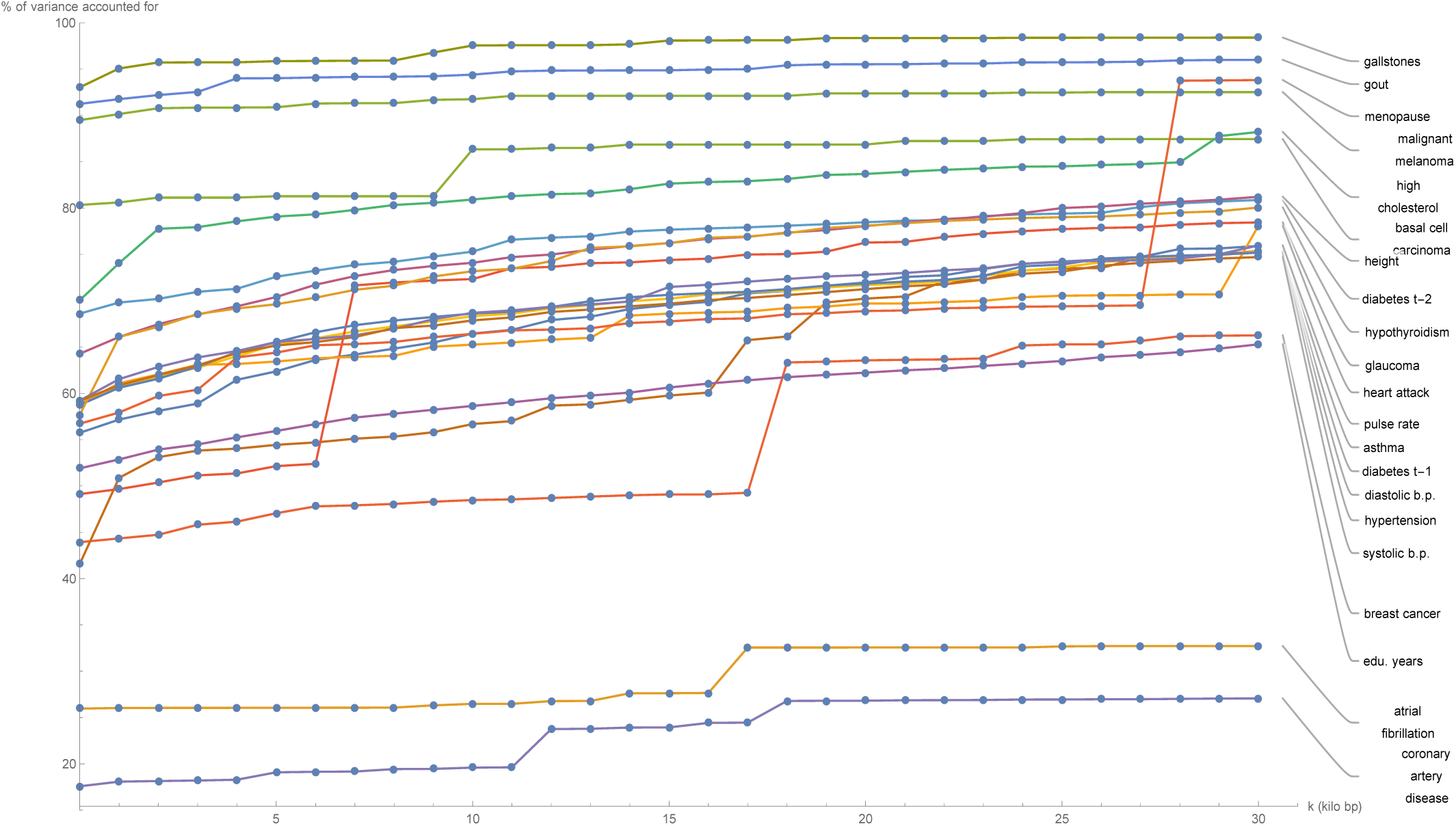
Plots of the variance accounted for by predictor SNPs located within genic regions, expressed as a percentage of the total variance accounted for by all predictor SNPs for that disease condition, against expansion of GENCODE Release 19 gene boundaries by k kilo base pairs..

Now that the extent of the variance accounted for by SNPs located within genic regions has been established for each disease condition, it is natural to investigate next how this genic variance is distributed between individual (protein-coding) genes. When considering a specific disease condition, for each gene, the variance accounted for by all predictor SNPs located within the (effective) gene boundary coordinates is summed and expressed as a percentage of the total variance accounted for by all predictor SNPs for that condition. For the purposes of this calculation, the boundaries of the genic regions were chosen to be at *k* = 30.

Figures 8 through 28 in Appendix A show the percentage of predictor variance accounted for by single genes for asthma, atrial fibrillation, basal cell carcinoma, breast cancer, coronary artery disease, type-1 diabetes, type-2 diabetes, diastolic blood pressure, educational years, gallstones, glaucoma, gout, heart attack, height, high cholesterol, hypertension, hypothyroidism, malignant melanoma, menopause, pulse rate, and systolic blood pressure. Only the fifteen largest values of variance accounted for by a single gene are displayed for each condition. As each genic predictor SNP may lie within the boundaries of more than one gene due to the expanded gene boundaries overlapping, multiple genes may share the exact same set of predictor SNPs and therefore the same value of total variance accounted for by single genes.

**Figure 3:**
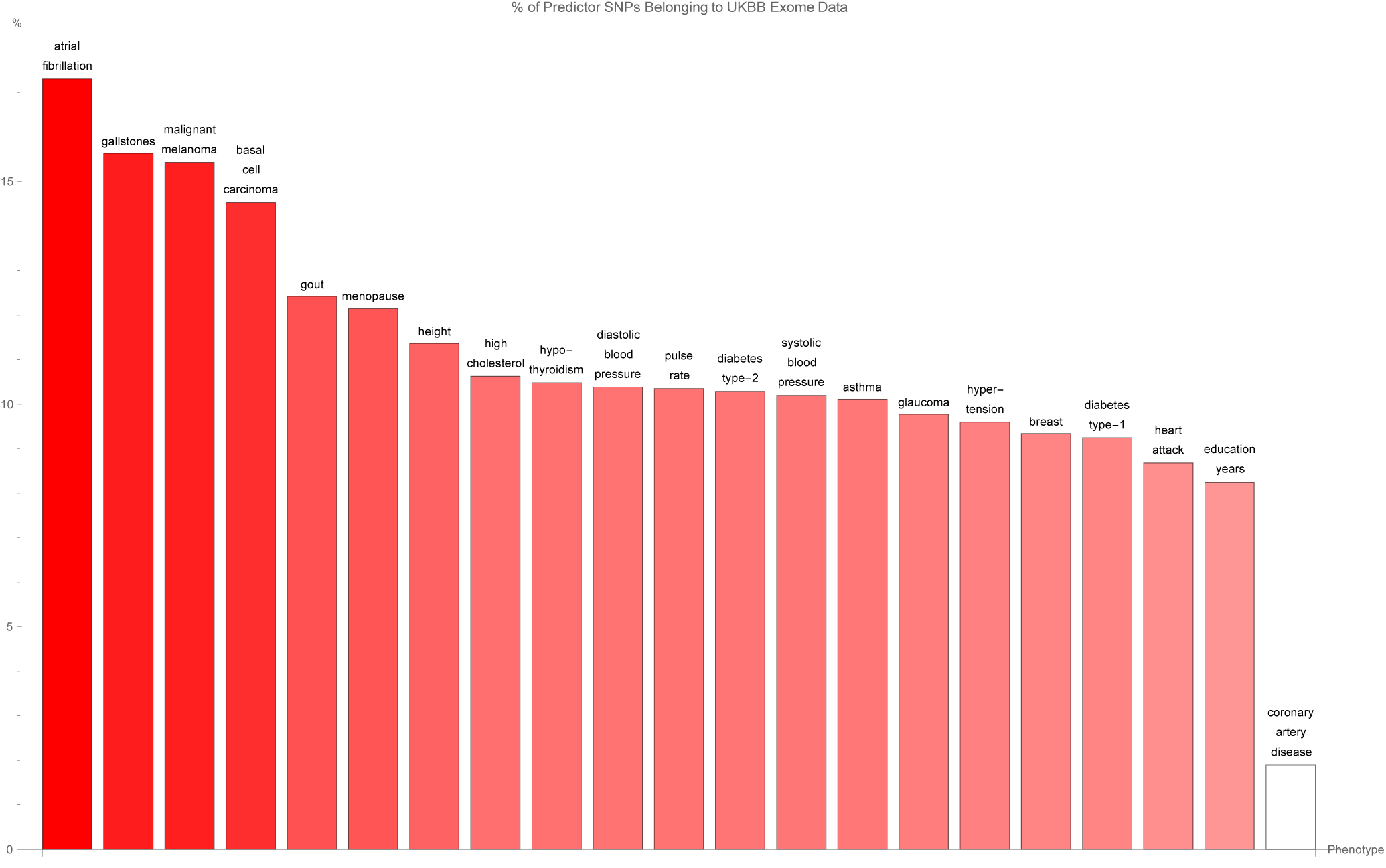
The percentage of predictor SNPs which are found both in genic regions and the UK Biobank exome data, for each disease condition. The disease conditions are listed from left to right on the horizontal axis in order of decreasing percentage. Each vertical bar is colored red with a depth of shade proportional to the height of the bar. Here, “genic” SNPs are contained within the GENCODE Release 19 gene boundaries plus 30 kilo base pairs at both ends.

**Figure 4:**
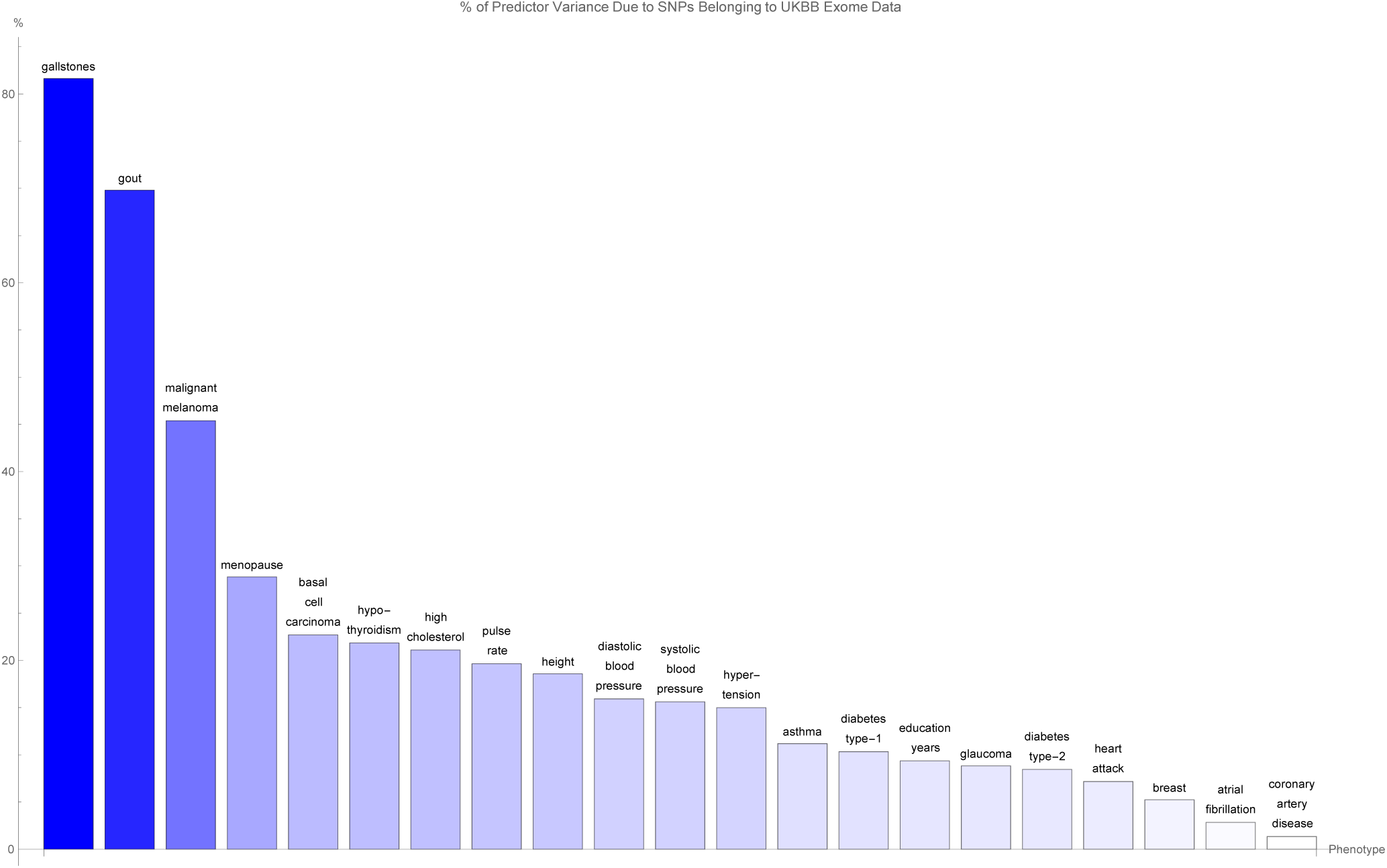
The variance accounted for by predictor SNPs which are both in genic regions and detected by the UK Biobank exome data, as a percentage of the total variance accounted for by all predictor SNPs, for each disease condition. The disease conditions are listed from left to right on the horizontal axis in order of decreasing percentage. Each vertical bar is colored blue with a depth of shade proportional to the height of the bar. Here, “genic” SNPs are contained within the GENCODE Release 19 gene boundaries plus 30 kilo base pairs at both ends.

**Figure 5:**
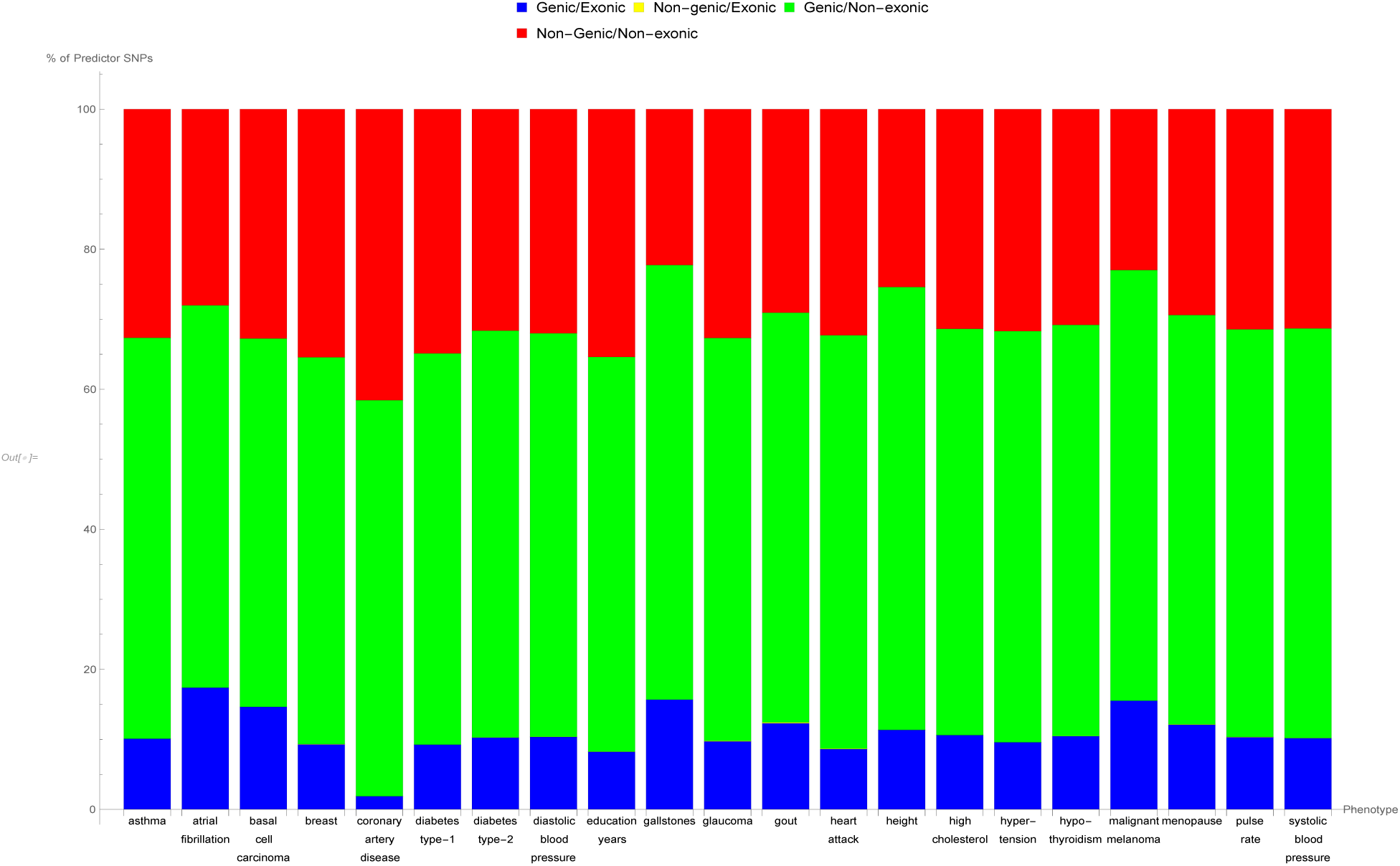
The breakdown of the percentage of predictor SNPs according to whether their location is genic and whether the exome data serves to probe them, for each predictor. The bar sections representing predictor SNPs in genic regions and in the exome data are labelled ‘Genic/Exonic’ and colored blue, those representing predictor SNPs not in genic regions but present in the exome data are ‘Non-genic/Exonic’ and colored yellow, those representing predictor SNPs which are located in genic regions but not found in the exome data are ‘Genic/Non-exonic’ and colored green, and those representing predictor SNPs neither in genic regions nor in the exome data are ‘Non-genic/Non-exonic’ and colored red. As expected, the yellow ‘Non-genic/Exonic’ bar sections are too small to be discernible. Here, “genic” SNPs are contained within the GENCODE Release 19 gene boundaries plus 30 kilo base pairs at both ends.

**Figure 6:**
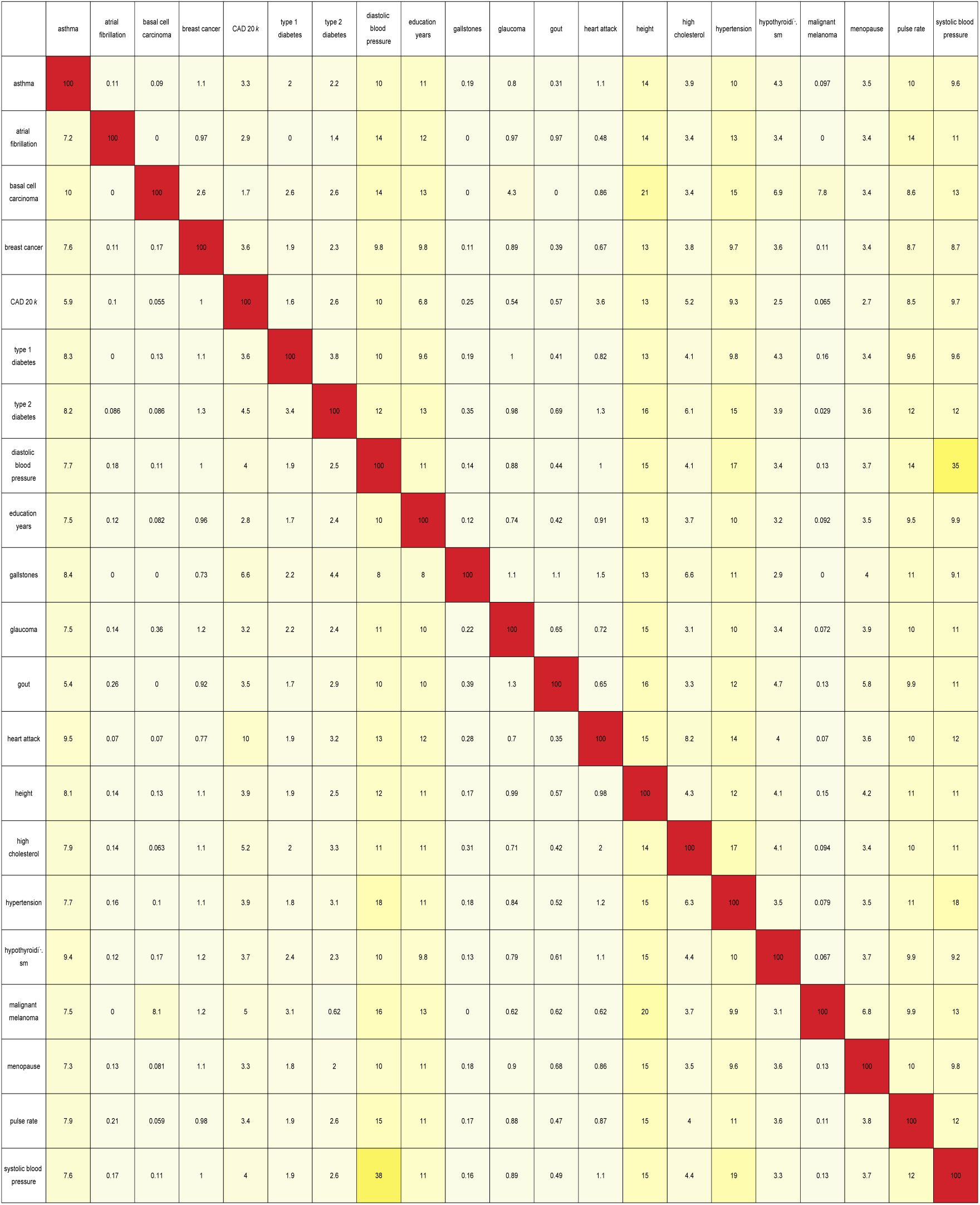
Table containing the pairwise overlap between predictors in terms of the number of predictor SNPs in common, expressed as a percentage of the total number of SNPs in the predictor for the condition labelling each row. Pairs of SNPs less than 4000 base pairs apart were considered to be identified with one another. For a particular row, the number in each column represents the percentage of SNPs from the row predictor that can be identified with the SNPs from the column predictor.

**Figure 7:**
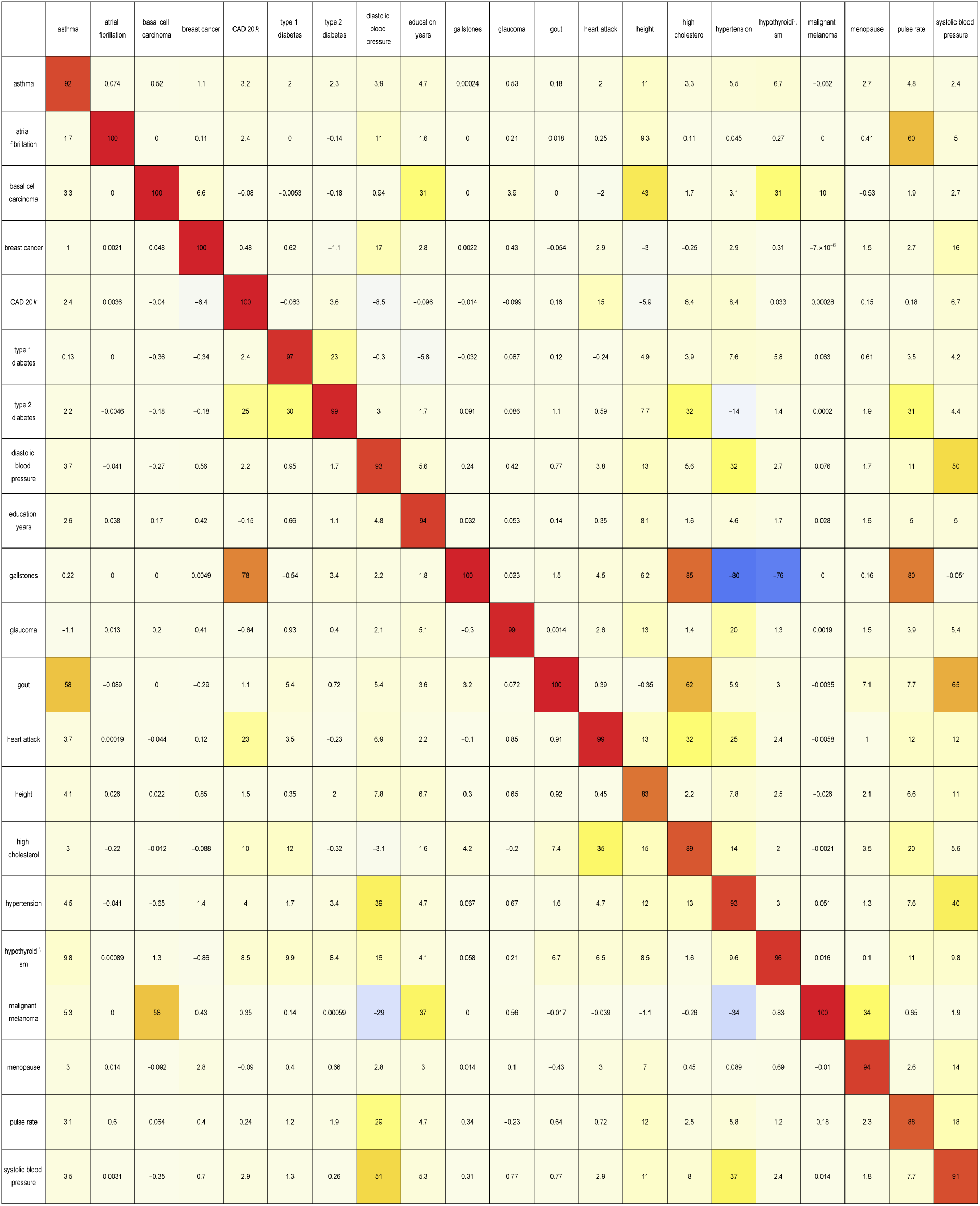
As described by Eq.(5), this table contains the pairwise overlap between predictors in terms of the variance accounted for by predictor SNPs in common, expressed as a percentage of the total variance accounted for by the SNPs belonging to the predictor for the condition labelling each row. The overlapping variance is weighted according to the sign of the correlation between each pair of associated SNPs 4000 kilo base pairs or less apart. This can cause diagonal elements to be less than one hundred percent, as anti-correlated SNPs may be included in the overlap calculation.

It is evident that certain disease conditions have one (or perhaps two or three) dominating value(s) of variance accounted for by a single gene. The most striking results are found with basal cell carcinoma (one gene, IRF4, supplying 28% of the predictor variance out of the 87% total genic variance at *k* = 30 and a second, TGM3, supplying 12% of the predictor variance), breast cancer (two genes, FGFR2 and TOX3, each supplying 14%-15% of the predictor variance out of 66% total genic variance), type-2 diabetes (one gene, TCF7L2, supplying 25% of the predictor variance out of 81% total genic variance), gallstones (three genes, ABCG8, ABCG5 and DYNC2LI1, each supplying 80%-83% of the predictor variance out of 98% total genic variance), glaucoma (two genes, ALDH9A1 and TMCO1, each supplying 19% of the predictor variance out of 78% total genic variance), gout (one gene, ABCG2, supplying 57% of the predictor variance out of 96% total genic variance and another, SLC2A9, supplying 12% of the predictor variance), heart attack (one gene, LPA, supplying 21% of the predictor variance out of 78% total genic variance), malignant melanoma (five genes, AC092143.1, TUBB3, TCF25, MC1R and DEF8, each supplying 37%-43% of the predictor variance out of 92% total genic variance), and menopause (one gene, UTY, supplying 24% of the predictor variance out of 94% total genic variance).

The more novel among these results are the strong links observed between glaucoma and ALDH9A1, between gallstones and DYNC2LI1, between menopause and UTY, and between malignant melanoma and the four genes AC092143.1, TUBB3, TCF25, and DEF8. These relationships are either not well-established up until now, or have not yet been suggested to be possible.

The rest of our findings confirm previous work by other groups. The association of IRF4 [44] and TGM3 with basal cell carcinoma is well-known, as is the relationship of FGFR2 [46] and TOX3 [47] to breast cancer, the link between TCF7L2 [48] and type-2 diabetes, the association of ABCG8 [49] and ABCG5 [50] with gallstones, the role of TMCO1 [51] in glaucoma, the relationship of ABCG2 [52, 53] and SLC2A9 [54] to gout, the connection between LPA [55] and heart attack / coronary artery disease, and the role of MC1R [56, 57] in malignant melanoma.

For every disease condition considered here, the full list of genes responsible for the top fifteen values of variance displayed here can be found in Appendix B.

### Overlap between predictor SNPs and whole-exome sequencing data

Modern whole-exome sequencing techniques are expected to be able to access about 85% of known disease-related variants [58, 59]. Assuming this is correct, we would expect about the same fraction of genic SNPs belonging to our predictors for disease conditions to be identifiable via exome-sequencing data. To verify this, we compared our sets of predictor SNPs against the whole-exome sequencing data released by the UK Biobank in March 2019 [60] (to be specific, the version of the whole-exome sequencing data set generated using a Functionally Equivalent (FE) pipeline [61]). In this section, once again, genic SNPs are defined as those SNPs located within the GENCODE Release 19 protein-coding gene boundaries extended by 30 kilo base pairs at both ends (or *k* = 30).

Figure 3 shows for each disease condition the percentage of predictor SNPs located in genic regions which are also found in the UK Biobank exome data, where the disease conditions are listed from left to right on the horizontal axis in order of decreasing percentage. For about two-thirds of the conditions surveyed, 10% or so of the genic SNPs for each predictor also formed part of the set of SNPs identified via exome-sequencing. The remaining disease conditions have up to about 17% (in terms of numbers) of their genic predictor SNPs detected by the exome-sequencing data set - with one exception, coronary artery disease, which displays an extraordinarily low (2%) value of this overlap.

Figure 4 shows for each disease condition the variance accounted for by the genic predictor SNPs from figure 3, expressed as a percentage of the total variance accounted for by all the SNPs in the predictor, where the disease conditions are listed from left to right on the horizontal axis in order of decreasing percentage. For the majority of conditions, 20% or less of the total variance accounted for by the predictor SNPs comes from genic SNPs which show up in the UK Biobank exome data. In fact, less than 5% of the variance accounted for by the the breast cancer, atrial fibrillation, and coronary artery disease predictor SNPs is detected by the exome data. Exceptions to this are the predictors for gallstones (80% of predictor variance detected by the exome data), gout (70% of predictor variance detected by the exome data), malignant melanoma (45% of predictor variance identified by the exome data), and menopause (30% of predictor variance identified by the exome data).

It may be worth noting that the ordering of the predictors by percentage of predictor SNPs which are genic (in Figure 1) and the ordering according to percentage of predictor SNPs which are both genic and accessible via the exome data in (Figure 3) is about the same. A similar observation can be made regarding Figures 2 and 4 (here in terms of variance accounted for).

In Figure 5, an overall view of the extent to which predictor SNPs in genic and non-genic regions are identifiable based on exome-sequencing data is given. Figure 5 shows for each disease condition the breakdown of the percentage of predictor SNPs according to whether their location is genic and whether the exome data serves to probe them. This breakdown does not vary much between predictors: in general about 10%-17% of predictor SNPs are both in genic regions and found in the exome data (‘Genic/Exonic’, colored blue), 0% (as might be expected) are not in genic regions but are present in the exome data (‘Non-genic/Exonic’, colored yellow), 55%-65% are located in genic regions but not found in the exome data (‘Genic/Non-exonic’, colored green), and 20%-35% are neither in genic regions nor in the exome data (‘Non-genic/Non-exonic’, colored red). The only deviation comes from the coronary artery disease predictor SNPs - less than 5% are ‘Genic/Exonic’, while nearly 45% are ‘Non-genic/Non-exonic’.

### Pairwise comparison of predictors

We now focus on finding connections between disease conditions based on similarities in their predictors. In this section, the analysis involving the coronary artery disease predictor was restricted to the top twenty thousand predictor SNPs as ranked by value of variance accounted for.

Figure 6 shows the pairwise overlap between disease conditions in terms of the number of predictor SNPs that each pair of conditions has in common. Here, pairs of SNPs less than 4000 base pairs apart were considered to be essentially the same SNP (where this separation was chosen so that most or all SNP pairs with high linkage disequilibrium levels are expected to be identified with one another [62]). Each row of the table corresponds to a particular disease condition, and reading the row from left to right (going from one column to the next) gives the number of predictor SNPs (expressed as a percentage of the total number of SNPs in the row-label predictor) that the conditions labelling each column share with the condition that the row corresponds to.

A pair of conditions may be considered to have a significant connection if the percentage of SNPs in common is substantial when read off the table both ways. Analyzing the table in this manner produces two notable groupings, with all the conditions in each group having large pairwise overlap.

1. Asthma – diastolic blood pressure – hypertension – systolic blood pressure – education years – height: The first four disease conditions form a combination which is not too surprising, but the same cannot be said about the last two traits. The table shows that all possible pairs taken from these six conditions overlap by about 10% or so, with the following exceptions which exceed this level by some way: 38% of the systolic blood pressure SNPs also belong to the diastolic blood pressure predictor, while 35% of the diastolic blood pressure SNPs also belong to the systolic blood pressure predictor; 17% of the diastolic blood pressure SNPs belong to the hypertension predictor, while 18% of the hypertension SNPs belong to the diastolic blood pressure predictor; and 18% of the hypertension SNPs belong to the systolic blood pressure predictor, while 19% of the systolic blood pressure SNPs belong to the hypertension predictor.
2. Basal cell carcinoma – malignant melanoma: 7.8% of the basal cell carcinoma SNPs belong to the malignant melanoma predictor, while 8.1% of the malignant melanoma SNPs belong to the basal cell carcinoma predictor.

Next, variance accounted for is used to measure the overlap, as seen in figure 7. This results in the discovery of more relations between different predictors. The method used on each pair of predictors was as follows: For each SNP, *i*, in the first predictor (corresponding to the condition labelling the row), all SNPs, *j*, from the second predictor (corresponding to the condition labelling the column) located less than 4000 base pairs away were identified. Every such associated SNP from the second predictor was assigned a weight of uniform magnitude, with the sign of the weight based on the sign of the SNP’s effect size relative to the sign of the effect size of the SNP from the first predictor - positive when the signs were the same, and negative when the signs were different. These weights were then multiplied by the variance due to the SNP from the first predictor and summed. If we label each set of SNPs within 4000 base pairs away as 𝒞_*i*_, this correlation estimate, 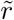 can be expressed as

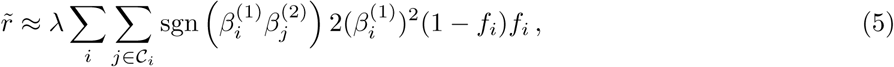

where the normalization is chosen to be the total predictor variance of the row predictor,

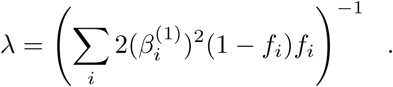

This produced a weighted overlap in terms of variance, that accounts for whether the associated SNP pairs are positively correlated (effect sizes have equal signs) or negatively correlated (effect sizes have differing signs). Figure 7 displays this overlap as a percentage, and is basically Figure 6 expressed in terms of variance (weighted according to correlation sign) accounted for by the overlapping predictor SNPs.

Once again, significant connections are observed in the case of diastolic blood pressure – hypertension – pulse rate – systolic blood pressure and basal cell carcinoma – malignant melanoma. In the case of the first group of conditions, the overlap of weighted variance ranges from 8% (systolic blood pressure – pulse rate) to 51% (systolic blood pressure – diastolic blood pressure). In the second case, 10% of the basal cell carcinoma predictor variance is shared with the malignant melanoma predictor, while 58% of the malignant melanoma predictor variance belongs to the basal cell carcinoma predictor.

We now also have groups of conditions which appear to be strongly positively correlated in terms of variance overlap, but were unremarkable in terms of number of SNPs in common. The most obvious is coronary artery disease – heart attack – high cholesterol – hypertension, where the magnitude of the pairwise overlap in terms of weighted variance ranges from 4% (hypertension – coronary artery disease) to 35% (high cholesterol – heart attack). Other single pairs of conditions with large overlapping variances are gout – high cholesterol (where 62% of the gout predictor variance also belongs to the high cholesterol predictor, while 35% of the high cholesterol predictor variance also belongs to the gout predictor), type-1 diabetes – type-2 diabetes (where 23% of the type-1 diabetes predictor variance also belongs to the type-2 diabetes predictor, while 30% of the type-2 diabetes predictor variance also belongs to the type-1 diabetes predictor), coronary artery disease – type-2 diabetes (4% and 25%), and gallstones – high cholesterol (85% and 4%). Also worth mentioning is the overlap of height with respectively: education years, asthma, diastolic blood pressure, hypertension, pulse rate, systolic blood pressure - which all are about 10%.

## Conclusions

This paper explores the genetic architectures of a number of common disease conditions and complex traits, as revealed by the most important SNPs used in genomic predictors.

The results are complex – primarily summarized in the many figures in the paper and Appendix. However, we can make some general statements:

I. The fraction of SNPs in or near genic regions varies widely by phenotype. For example, in the case of Coronary Artery Disease and Atrial Fibrillation, less than 20-30 percent of the total risk variance is due to SNPs near genic regions.
II. For the majority of disease conditions studied, *most* of the variance is accounted for by SNPs whose state cannot be determined from exome-sequencing data. This suggests that exome data alone will miss much of the heritability for these traits. Stated somewhat differently: exome sequencing data for a specific individual misses much of the information necessary to compute their PRS score!
III. The DNA regions used in disease risk predictors so far constructed seem to be largely disjoint (with a few interesting exceptions), suggesting that individual genetic disease risks are largely uncorrelated.

Observation III has interesting implications for pleiotropy [63–65]. We found that genetic risks are largely uncorrelated for different conditions. This suggests that there can exist individuals with, e.g., low risk simultaneously in each of multiple conditions, for any essentially any combination of conditions. There is no trade-off required between different disease risks (at least, not among the ones studied here). One could speculate that a lucky individual with exceptionally low risk across multiple conditions might have an unusually long life expectancy.

Of course, the same applies for high risk: some unlucky individuals have high risk for multiple conditions simultaneously. In fact, there appear to be combinations of SNPs that could make a specific individual an outlier in each of the conditions studied, simultaneously.

## Note

Recently, it was pointed out [66] that the processing of the whole-exome sequencing data via the FE pipeline had been carried out in a manner that failed to take into account the presence of alternative contigs in the GRCh38 reference genome. This is expected to have led to fewer variants being called than there should be in the resultant data set. Out of the total of 204,829 genomic regions comprising 39.20 MBp of the human genome targeted by the whole-exome sequencing process, data from 7554 regions extending across 1.53 MBp were potentially affected by this error.

In an analysis of this issue [67], Jia et al. compared the number of exome variants per gene identified when using whole-exome sequencing data from the UK Biobank versus using data from gnomAD. They found 641 genes for which the UK Biobank exome data contains no variants whatsoever. In contrast, they calculated that it is highly probable for the UK Biobank exome data to identify at least one variant per gene in the case of the vast majority (93%) of these 641 genes.

With the aim of gauging the extent to which our results may have been impacted by this discrepancy, we examined the overlap between our lists of top genes ranked by variance accounted for by predictor SNPs (Figures 29 - 49) and the 641 potentially problematic genes. We found that for most (17 out of 21) conditions, the genes responsible for the top fifteen values of variance accounted for do not include any of the 641 potentially problematic genes.

The exceptions to this are asthma, basal cell carcinoma, type-1 diabetes, and hypothyroidism. To be specific: Asthma has 2 genes (HLA-DQB1 and HLA-DQA1 with variance 2%-3% each) out of its top 18 genes included among the 641 potentially problematic genes. For comparison, asthma has 3% as the highest percentage of predictor variance accounted for by a single gene. Basal cell carcinoma has 1 gene (HLA-DQA1 with variance 2%) out of its top 20 genes included among the 641 potentially problematic genes, while 28% is the highest fraction of predictor variance accounted for by a single gene. Type-1 diabetes has 12 genes (HSPA1L, HLA-DRB1, BTNL2, HLA-DQB1, HLA-DQB2, NEU1, HLA-DOA, HLA-DQA1, HLA-DRA, HSPA1B, C6orf48, LSM2) out of its top 25 genes among the 641 potentially problematic genes. These 12 genes include the top four genes ranked by variance accounted for, with 14%, 9%, 7%, and 4% of predictor variance respectively. Hypothyroidism has 4 genes (HLA-DPA1, HLA-DQB1, HLA-DQA1, and HLA-DPB1, where two have 5% of predictor variance each and two have 1% each) out of its top 17 genes included among the 641 potentially problematic genes, while 8% is the highest value of predictor variance accounted for by a single gene.

Clearly, for all conditions except type-1 diabetes, the 641 potentially problematic genes play very little part in determining the variance accounted for by the predictor SNPs. We feel that this justifies our opinion that the upcoming corrections to the UK Biobank exome data will not qualitatively change our findings as regards these conditions. There is a possibility, of course, that there could be a significant shift in our results for the type-1 diabetes predictor.

## Acknowledgements

SY, TR, LL, and SH acknowledge support from the Office of the Vice-President for Research at MSU. LL was also supported by Genomic Prediction, Inc. during part of this project. This work was supported in part by Michigan State University through computational resources provided by the Institute for Cyber-Enabled Research. This research was conducted using the UK Biobank Resource under UK Biobank Main Application 15326.

## Appendix A

**Figure 8:**
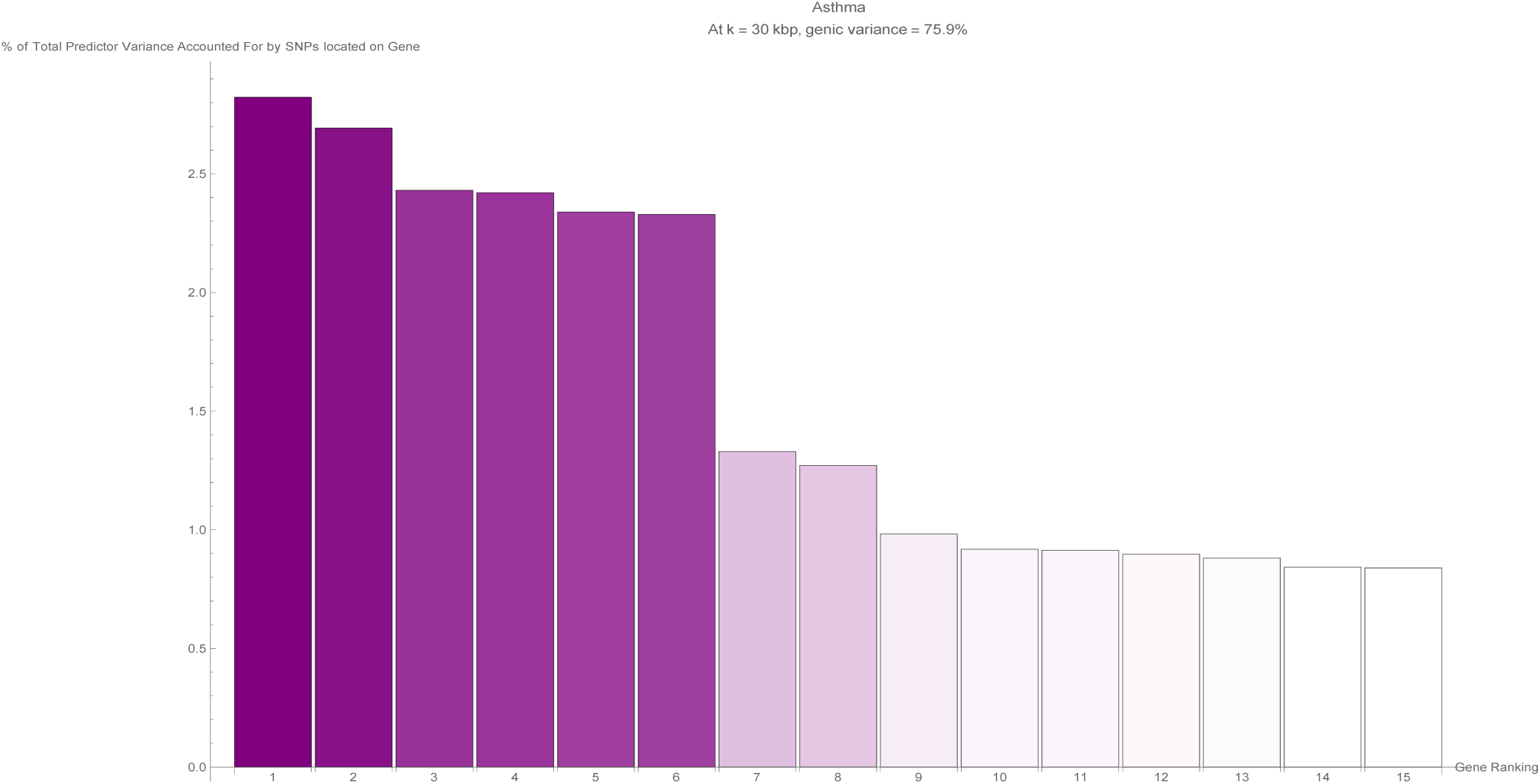
The fifteen largest total values of variance accounted for by predictor SNPs located on a single gene (in terms of the percentage of total variance accounted for by all predictor SNPs) for the asthma predictor. Each vertical bar is colored violet with a depth of shade proportional to the height of the bar. Here, ‘genic’ SNPs are contained within the GENCODE Release 19 gene boundaries plus 30 kilo base pairs at both ends.

**Figure 9:**
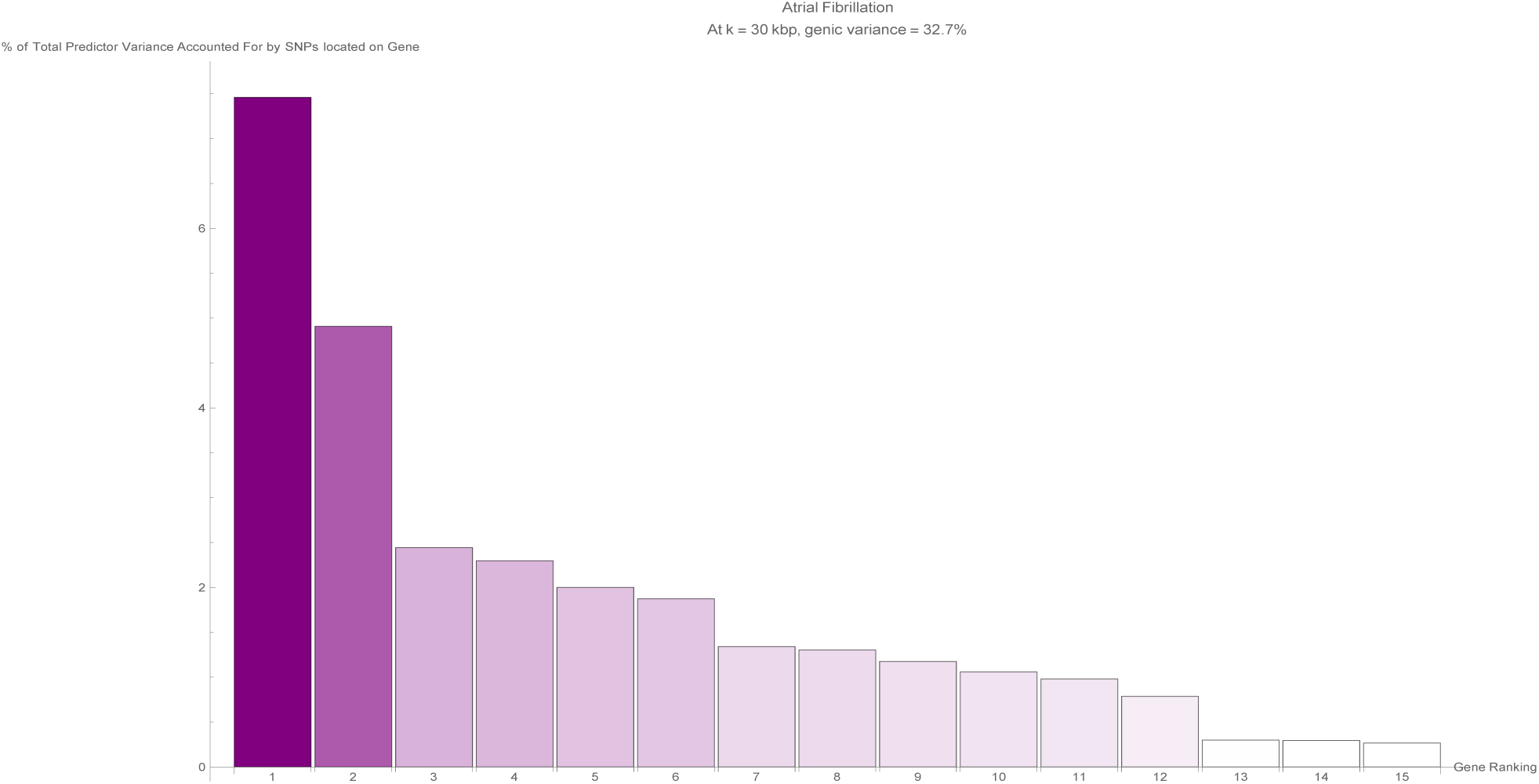
The fifteen largest total values of variance accounted for by predictor SNPs located on a single gene (in terms of the percentage of total variance accounted for by all predictor SNPs) for the atrial fibrillation predictor. Each vertical bar is colored violet with a depth of shade proportional to the height of the bar. Here, ‘genic’ SNPs are contained within the GENCODE Release 19 gene boundaries plus 30 kilo base pairs at both ends.

**Figure 10:**
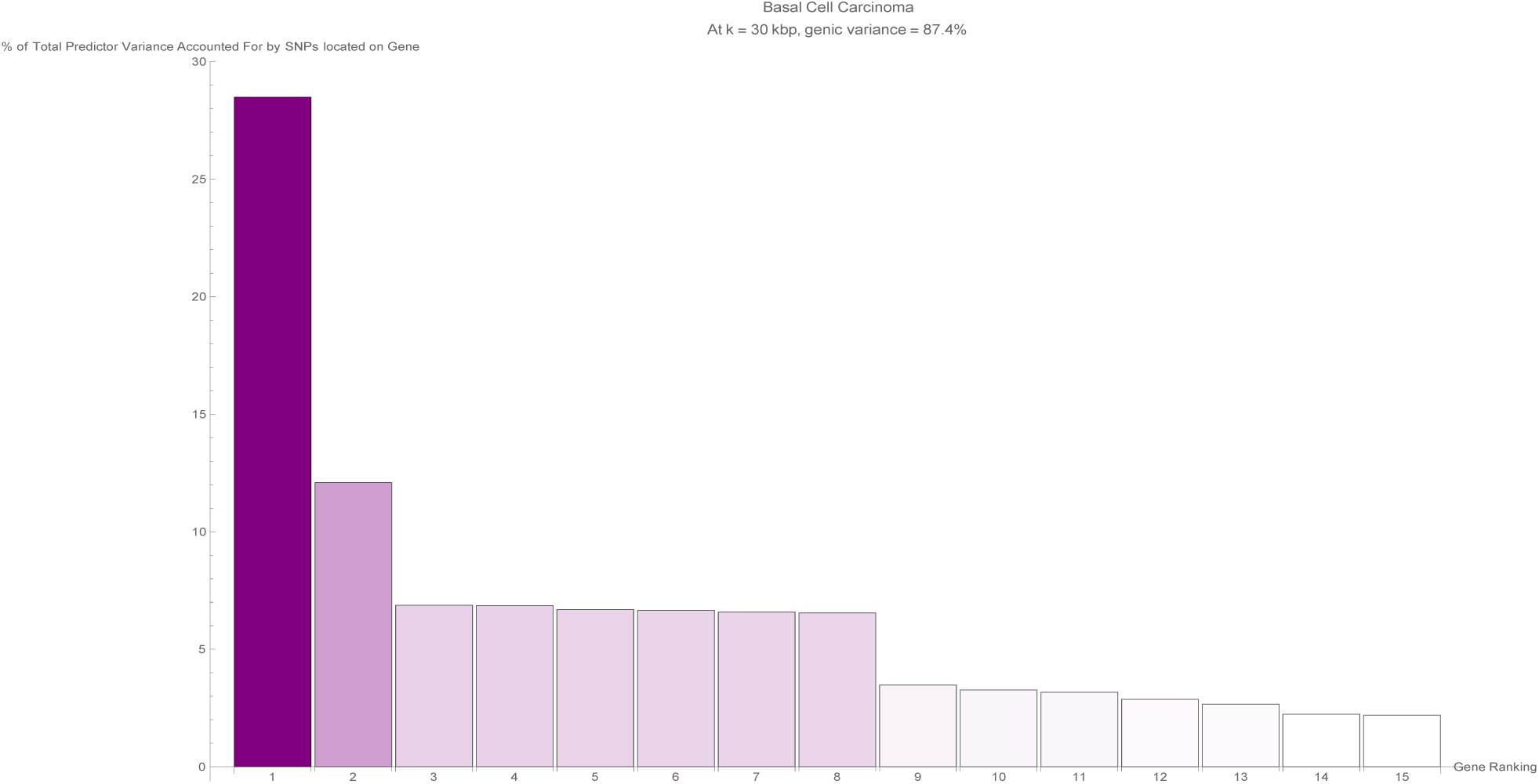
The fifteen largest total values of variance accounted for by predictor SNPs located on a single gene (in terms of the percentage of total variance accounted for by all predictor SNPs) for the basal cell carcinoma predictor. Each vertical bar is colored violet with a depth of shade proportional to the height of the bar. Here, ‘genic’ SNPs are contained within the GENCODE Release 19 gene boundaries plus 30 kilo base pairs at both ends.

**Figure 11:**
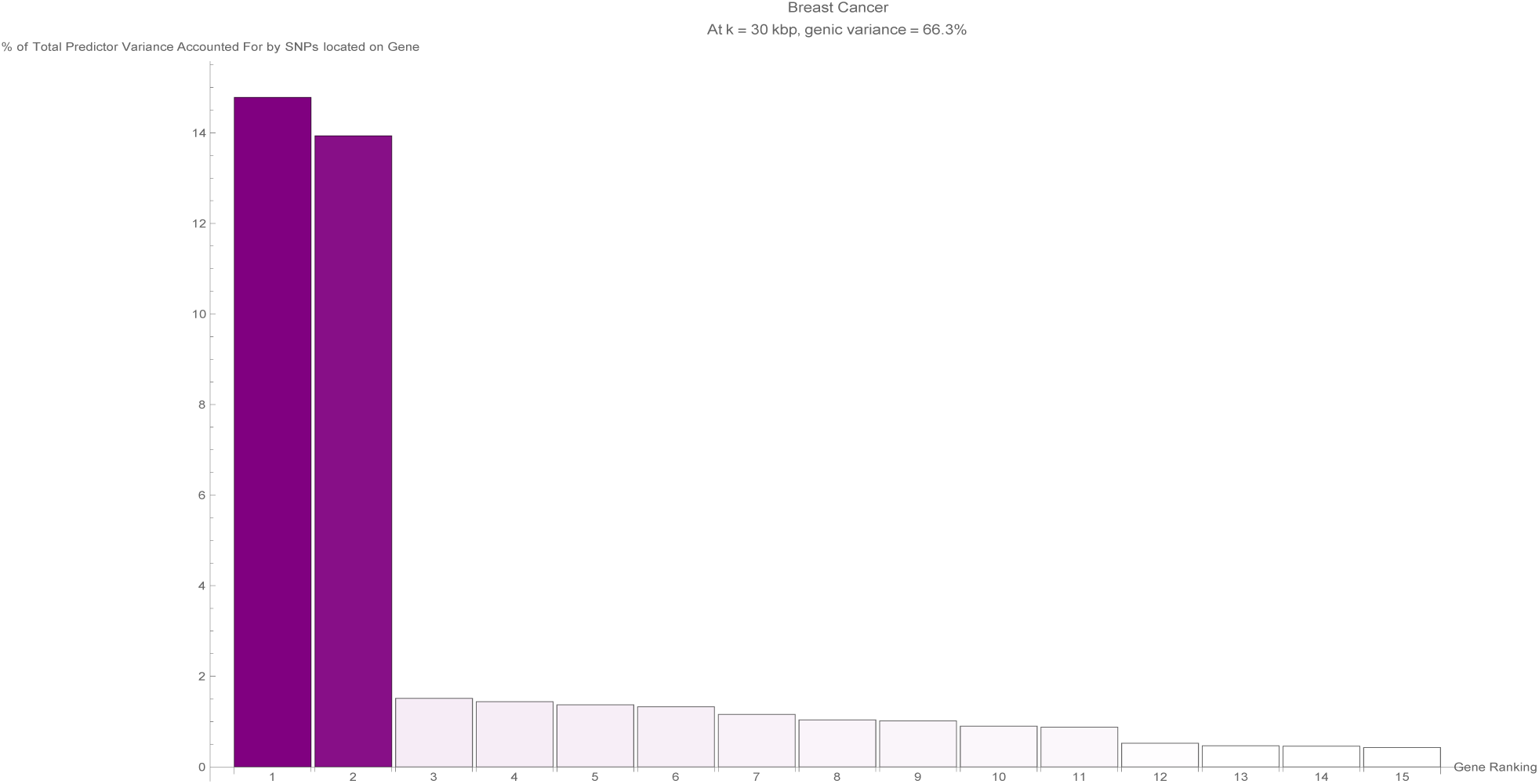
The fifteen largest total values of variance accounted for by predictor SNPs located on a single gene (in terms of the percentage of total variance accounted for by all predictor SNPs) for the breast cancer predictor. Each vertical bar is colored violet with a depth of shade proportional to the height of the bar. Here, ‘genic’ SNPs are contained within the GENCODE Release 19 gene boundaries plus 30 kilo base pairs at both ends.

**Figure 12:**
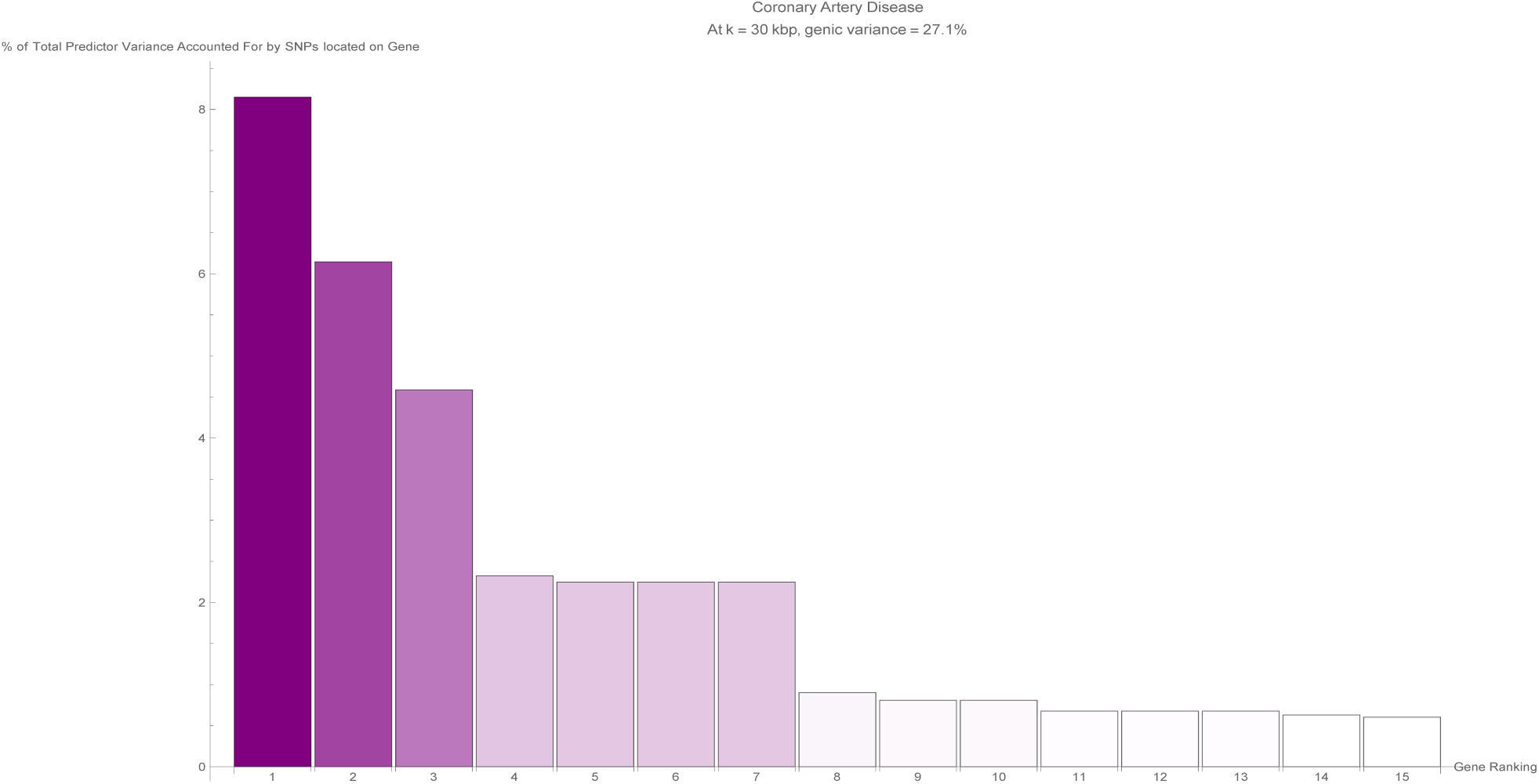
The fifteen largest total values of variance accounted for by predictor SNPs located on a single gene (in terms of the percentage of total variance accounted for by all predictor SNPs) for the coronary artery disease predictor. Each vertical bar is colored violet with a depth of shade proportional to the height of the bar. Here, ‘genic’ SNPs are contained within the GENCODE Release 19 gene boundaries plus 30 kilo base pairs at both ends.

**Figure 13:**
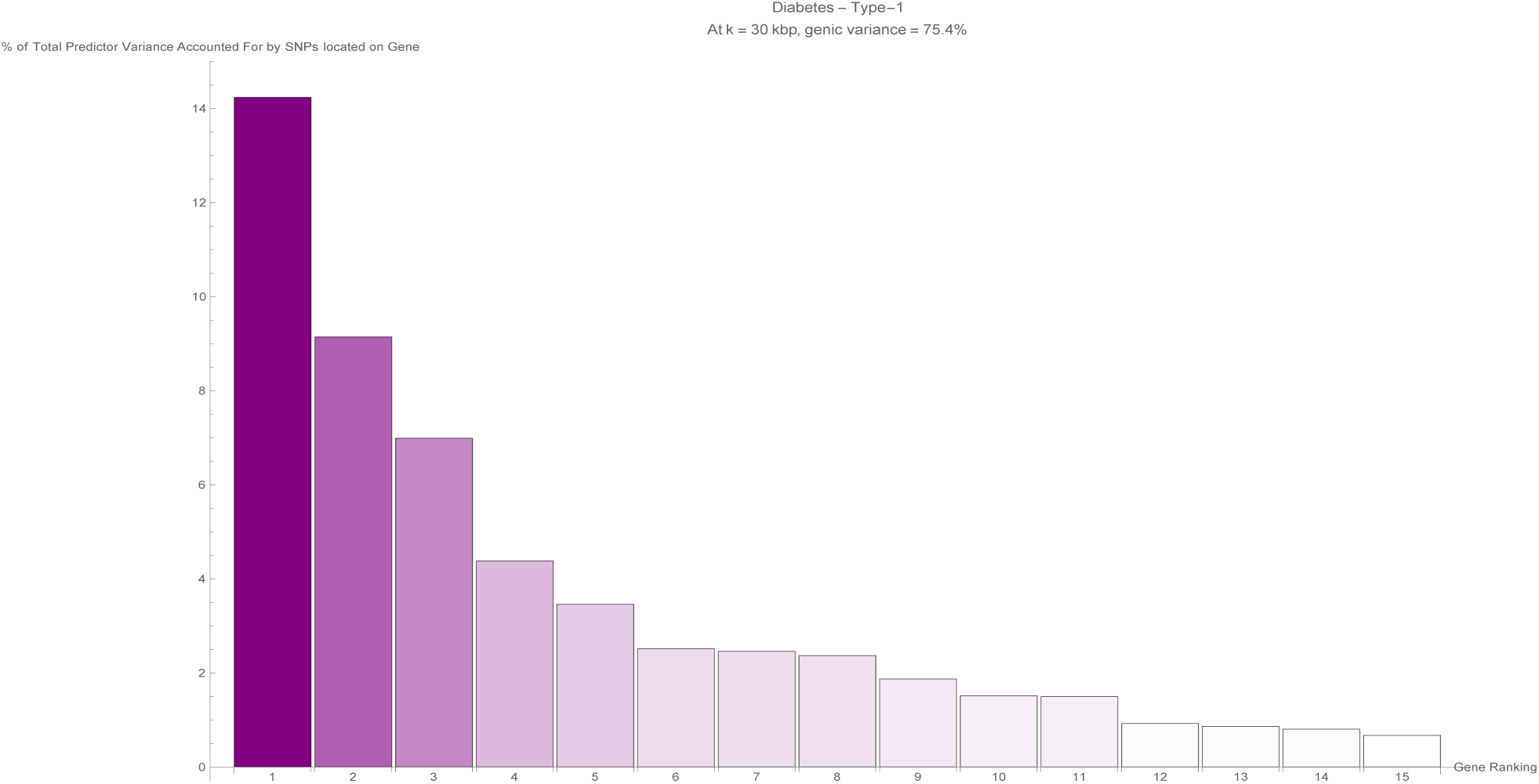
The fifteen largest total values of variance accounted for by predictor SNPs located on a single gene (in terms of the percentage of total variance accounted for by all predictor SNPs) for the type-1 diabetes predictor. Each vertical bar is colored violet with a depth of shade proportional to the height of the bar. Here, ‘genic’ SNPs are contained within the GENCODE Release 19 gene boundaries plus 30 kilo base pairs at both ends.

**Figure 14:**
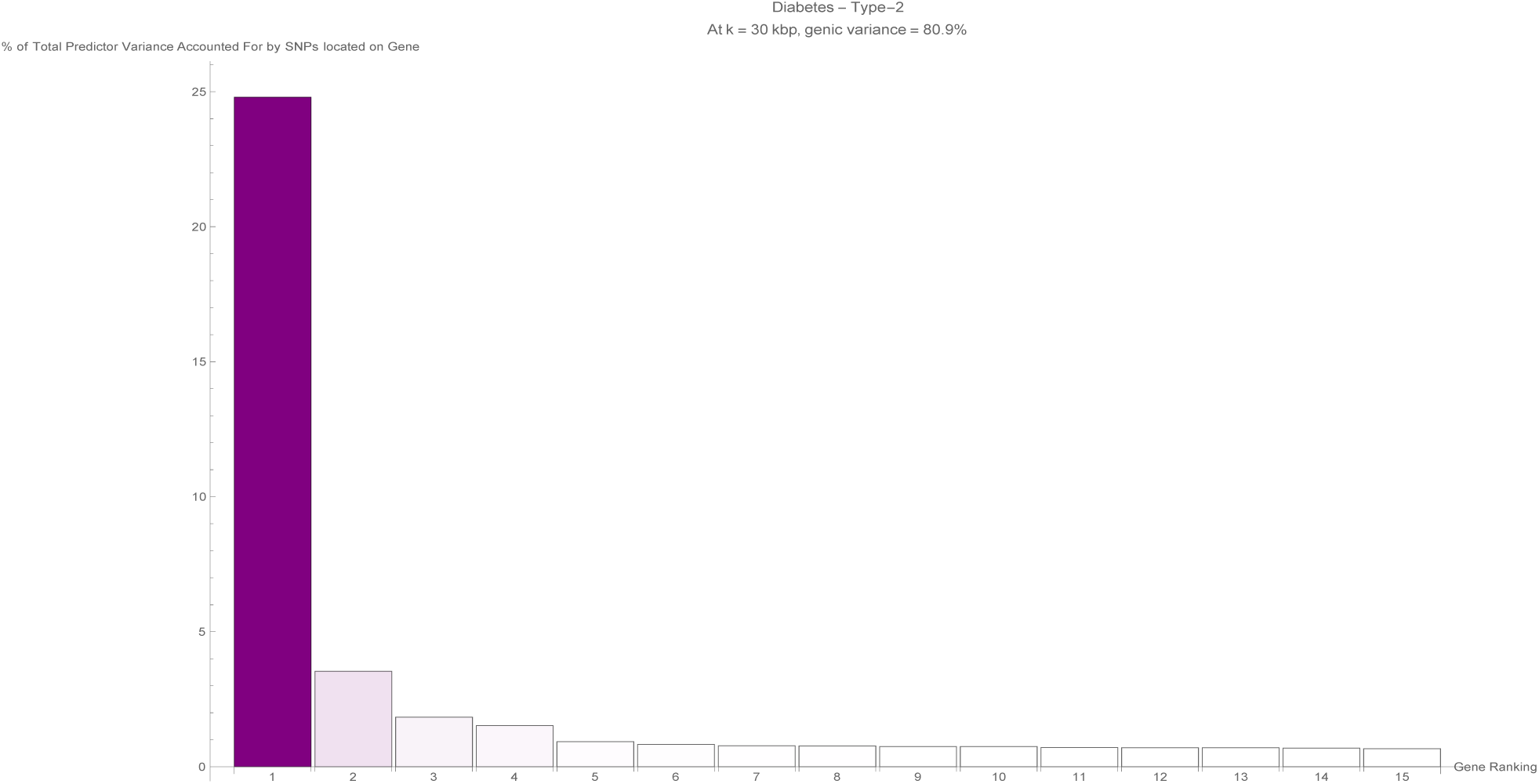
The fifteen largest total values of variance accounted for by predictor SNPs located on a single gene (in terms of the percentage of total variance accounted for by all predictor SNPs) for the type-2 diabetes predictor. Each vertical bar is colored violet with a depth of shade proportional to the height of the bar. Here, ‘genic’ SNPs are contained within the GENCODE Release 19 gene boundaries plus 30 kilo base pairs at both ends.

**Figure 15:**
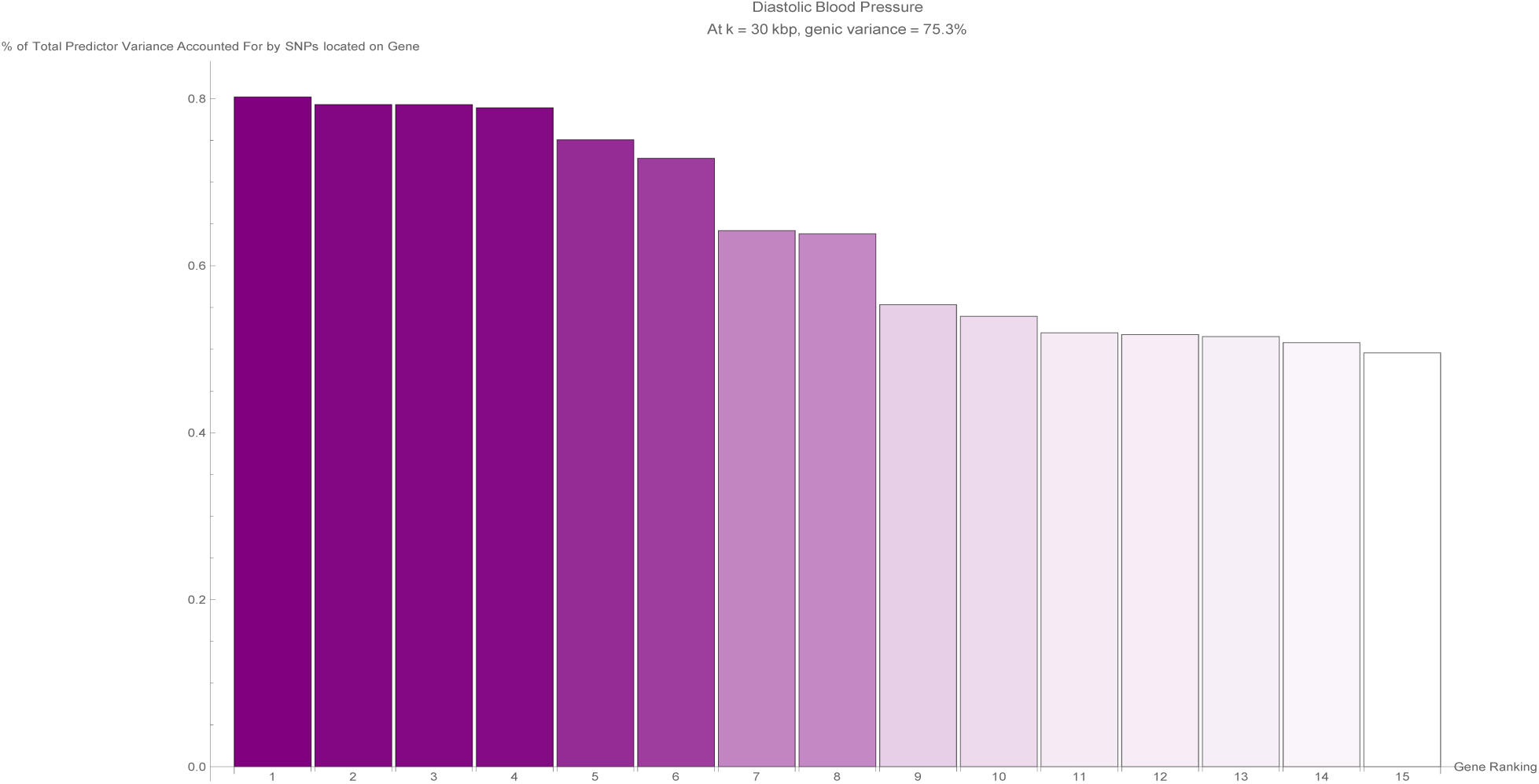
The fifteen largest total values of variance accounted for by predictor SNPs located on a single gene (in terms of the percentage of total variance accounted for by all predictor SNPs) for the diastolic blood pressure predictor. Each vertical bar is colored violet with a depth of shade proportional to the height of the bar. Here, ‘genic’ SNPs are contained within the GENCODE Release 19 gene boundaries plus 30 kilo base pairs at both ends.

**Figure 16:**
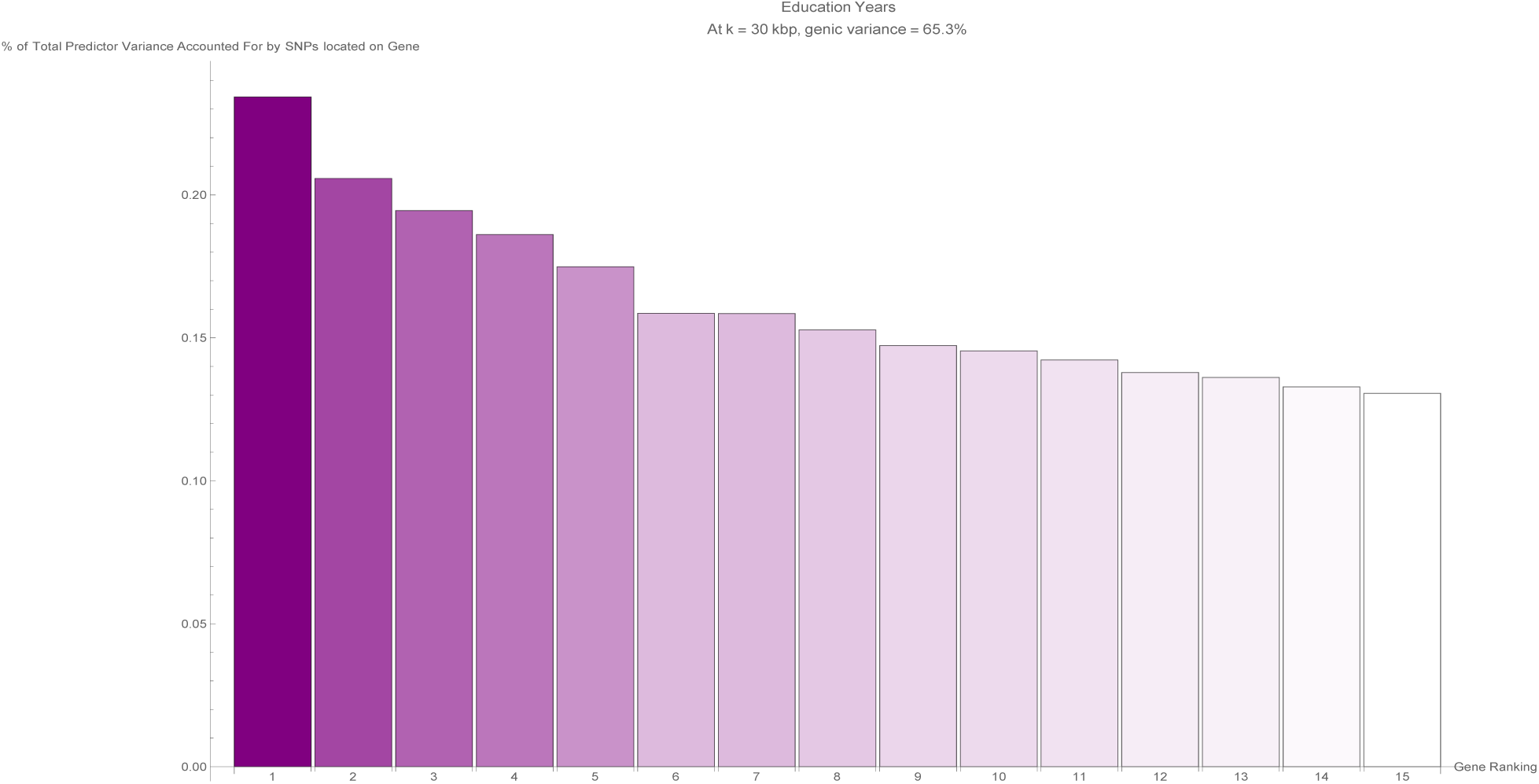
The fifteen largest total values of variance accounted for by predictor SNPs located on a single gene (in terms of the percentage of total variance accounted for by all predictor SNPs) for the education years predictor. Each vertical bar is colored violet with a depth of shade proportional to the height of the bar. Here, ‘genic’ SNPs are contained within the GENCODE Release 19 gene boundaries plus 30 kilo base pairs at both ends.

**Figure 17:**
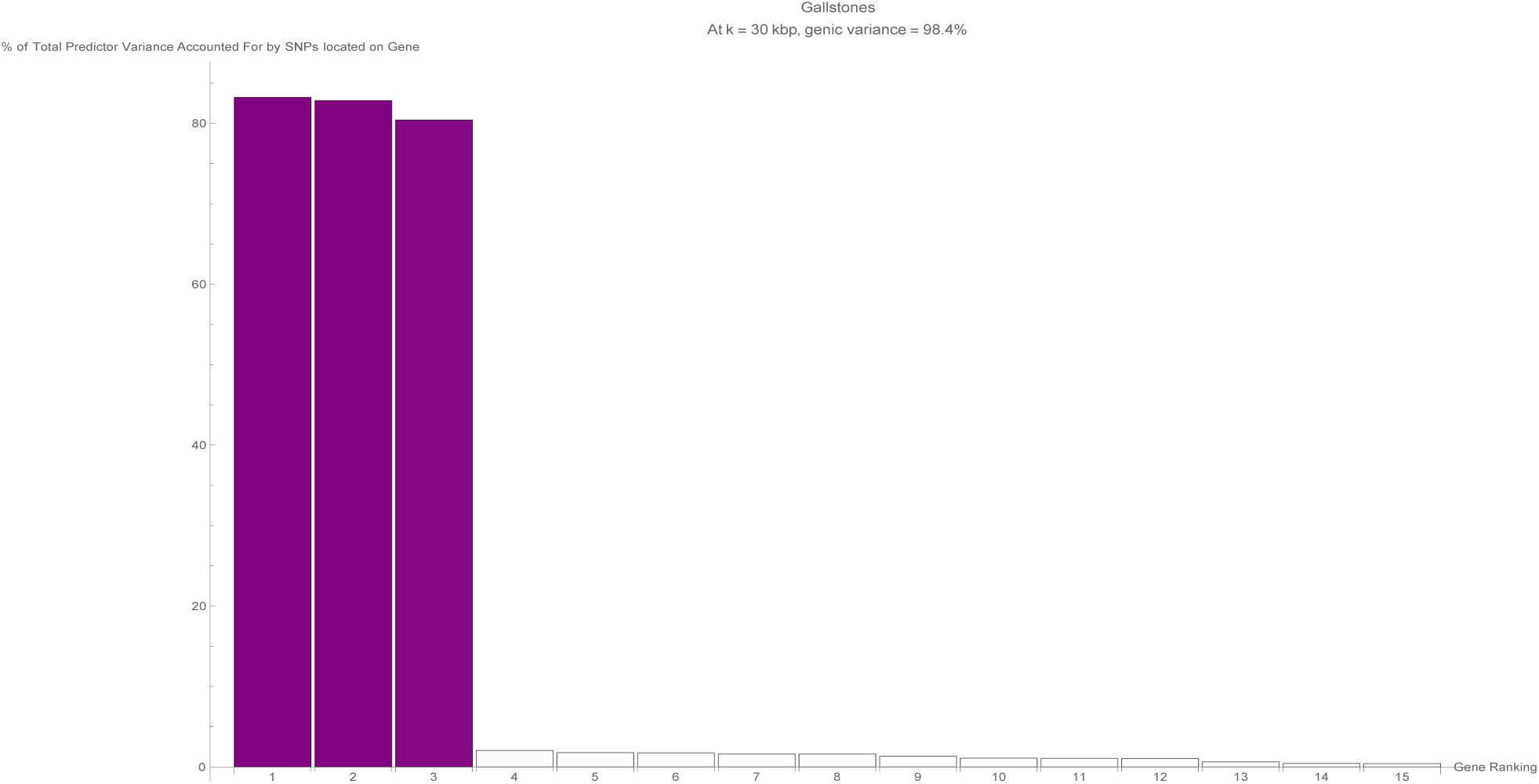
The fifteen largest total values of variance accounted for by predictor SNPs located on a single gene (in terms of the percentage of total variance accounted for by all predictor SNPs) for the gallstones predictor. Each vertical bar is colored violet with a depth of shade proportional to the height of the bar. Here, ‘genic’ SNPs are contained within the GENCODE Release 19 gene boundaries plus 30 kilo base pairs at both ends.

**Figure 18:**
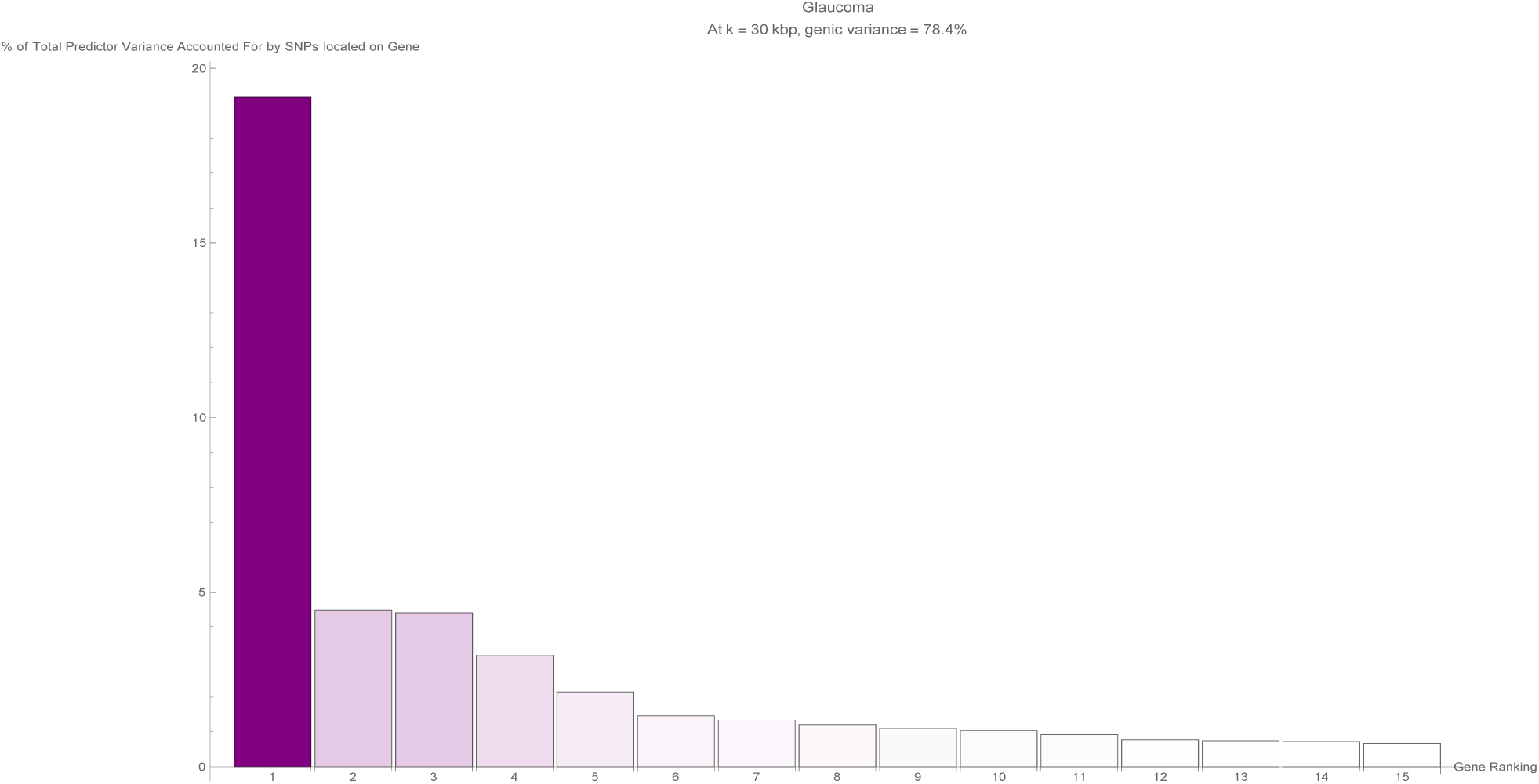
The fifteen largest total values of variance accounted for by predictor SNPs located on a single gene (in terms of the percentage of total variance accounted for by all predictor SNPs) for the glaucoma predictor. Each vertical bar is colored violet with a depth of shade proportional to the height of the bar. Here, ‘genic’ SNPs are contained within the GENCODE Release 19 gene boundaries plus 30 kilo base pairs at both ends.

**Figure 19:**
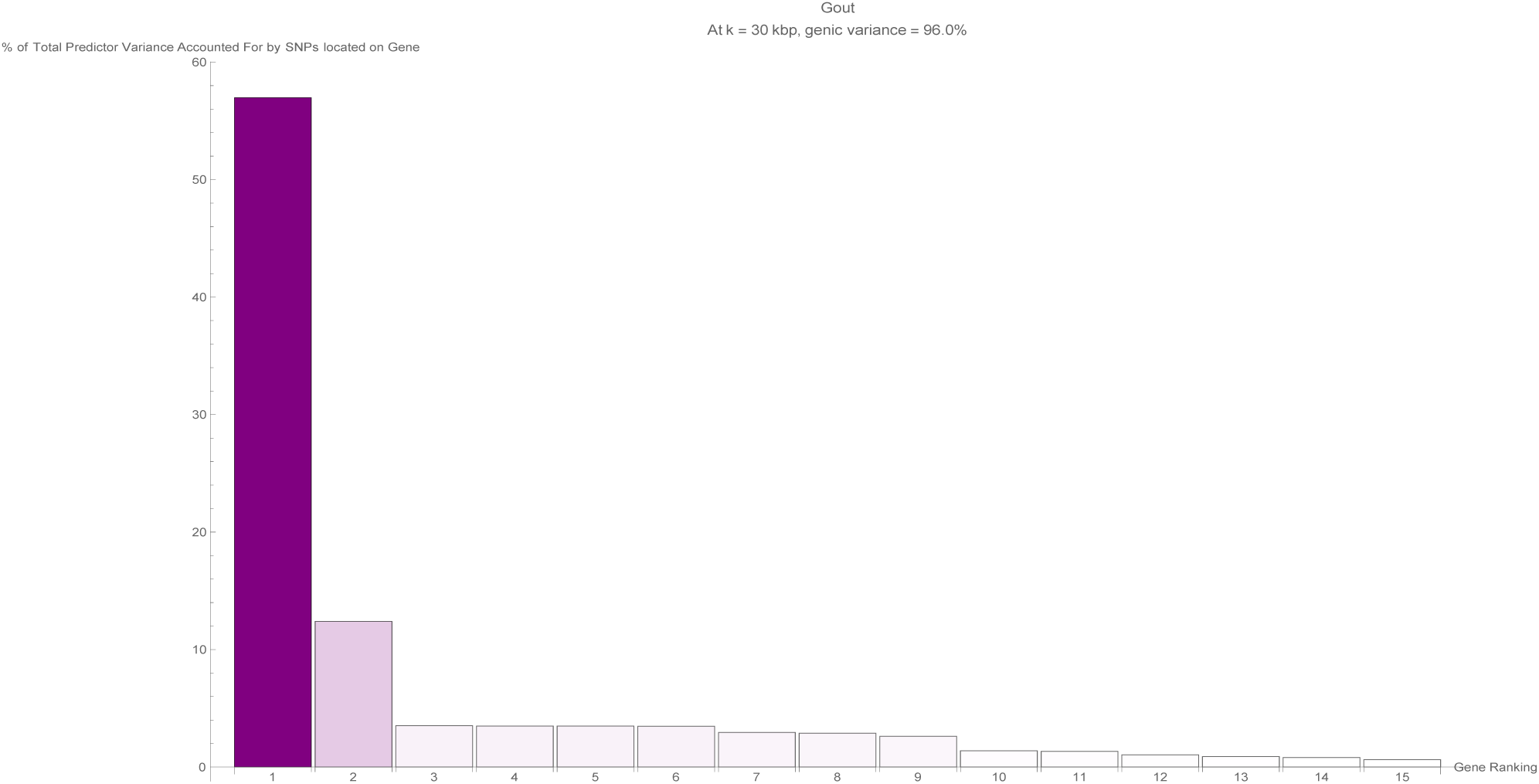
The fifteen largest total values of variance accounted for by predictor SNPs located on a single gene (in terms of the percentage of total variance accounted for by all predictor SNPs) for the gout predictor. Each vertical bar is colored violet with a depth of shade proportional to the height of the bar. Here, ‘genic’ SNPs are contained within the GENCODE Release 19 gene boundaries plus 30 kilo base pairs at both ends.

**Figure 20:**
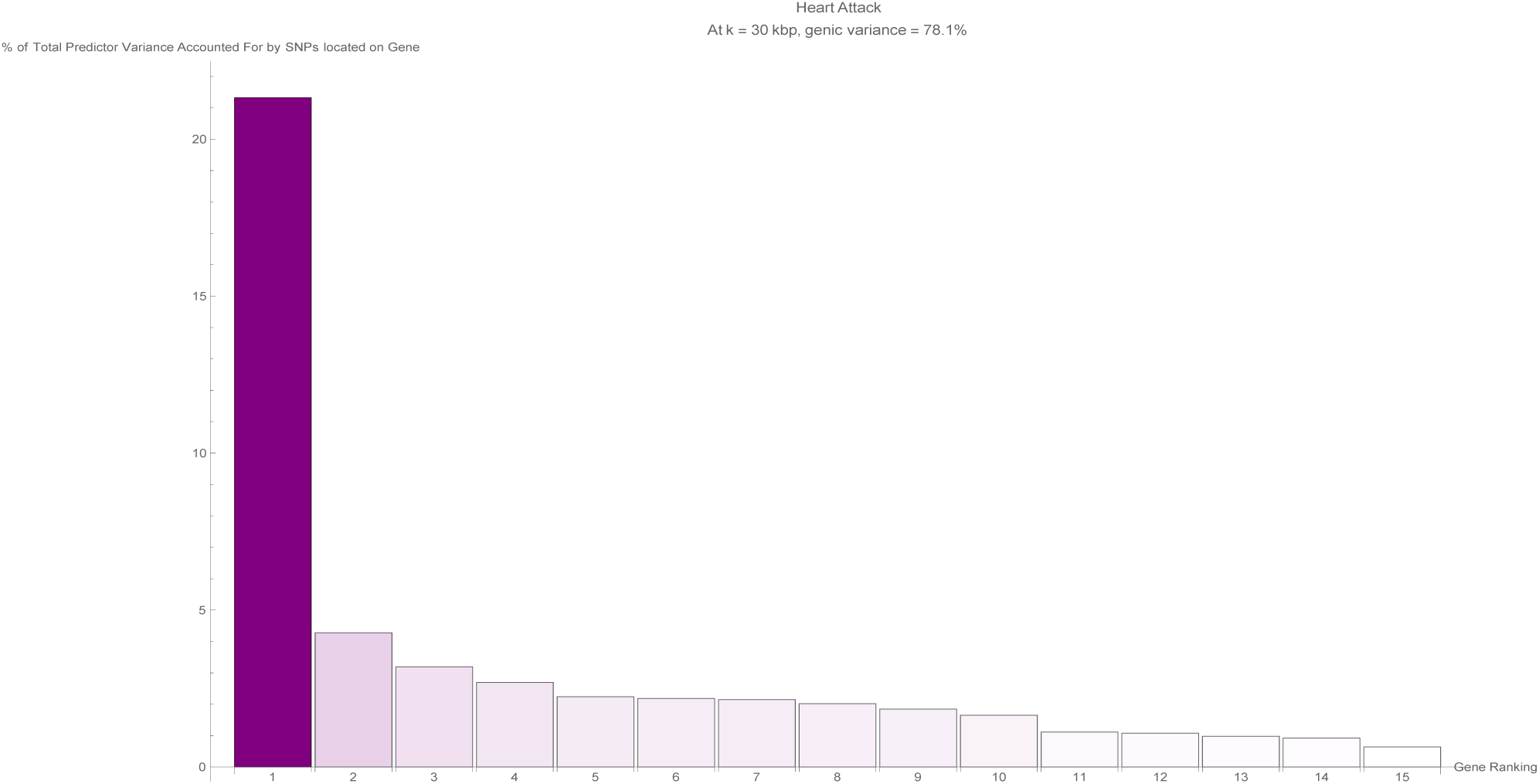
The fifteen largest total values of variance accounted for by predictor SNPs located on a single gene (in terms of the percentage of total variance accounted for by all predictor SNPs) for the heart attack predictor. Each vertical bar is colored violet with a depth of shade proportional to the height of the bar. Here, ‘genic’ SNPs are contained within the GENCODE Release 19 gene boundaries plus 30 kilo base pairs at both ends.

**Figure 21:**
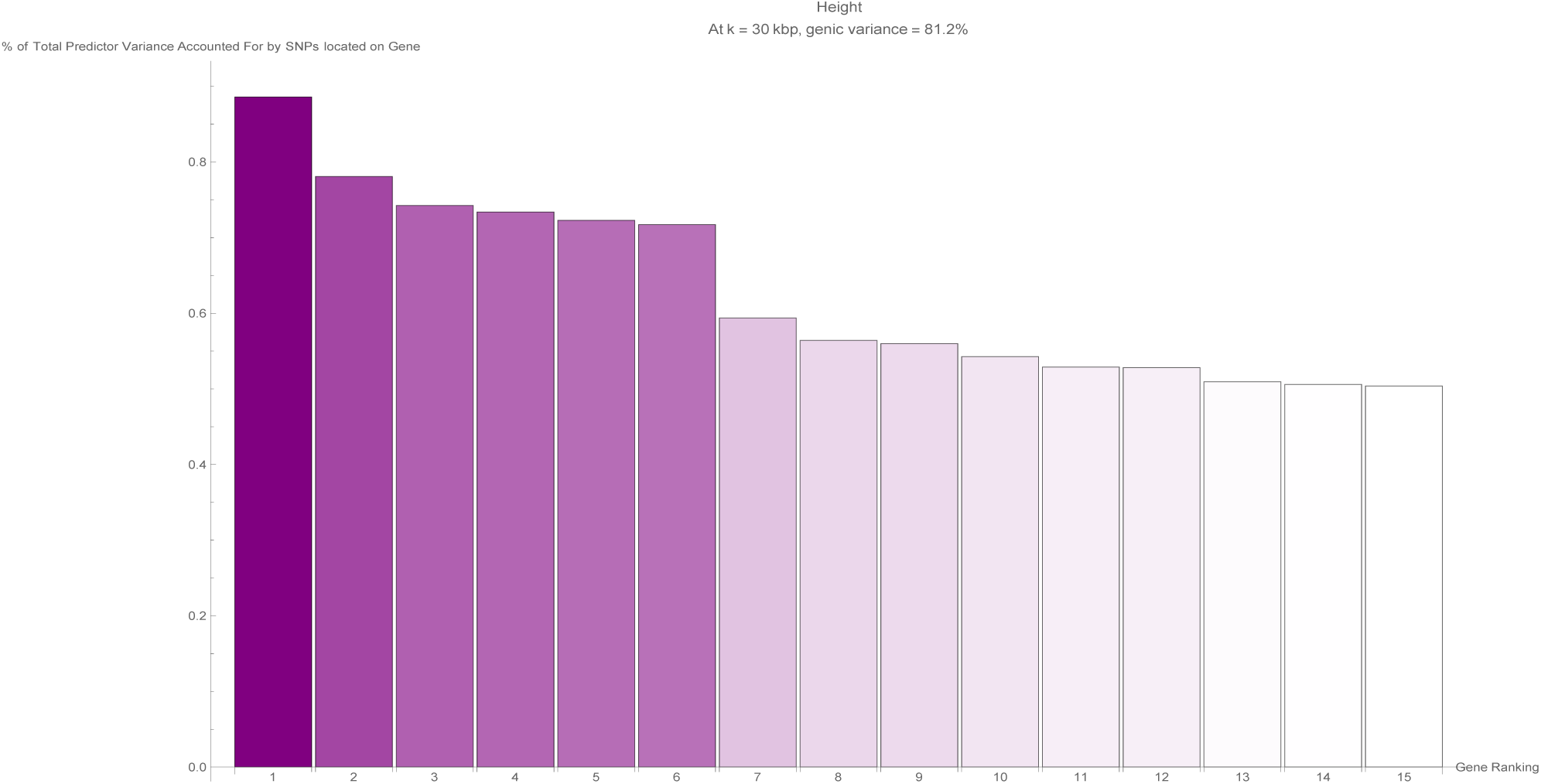
The fifteen largest total values of variance accounted for by predictor SNPs located on a single gene (in terms of the percentage of total variance accounted for by all predictor SNPs) for the height predictor. Each vertical bar is colored violet with a depth of shade proportional to the height of the bar. Here, ‘genic’ SNPs are contained within the GENCODE Release 19 gene boundaries plus 30 kilo base pairs at both ends.

**Figure 22:**
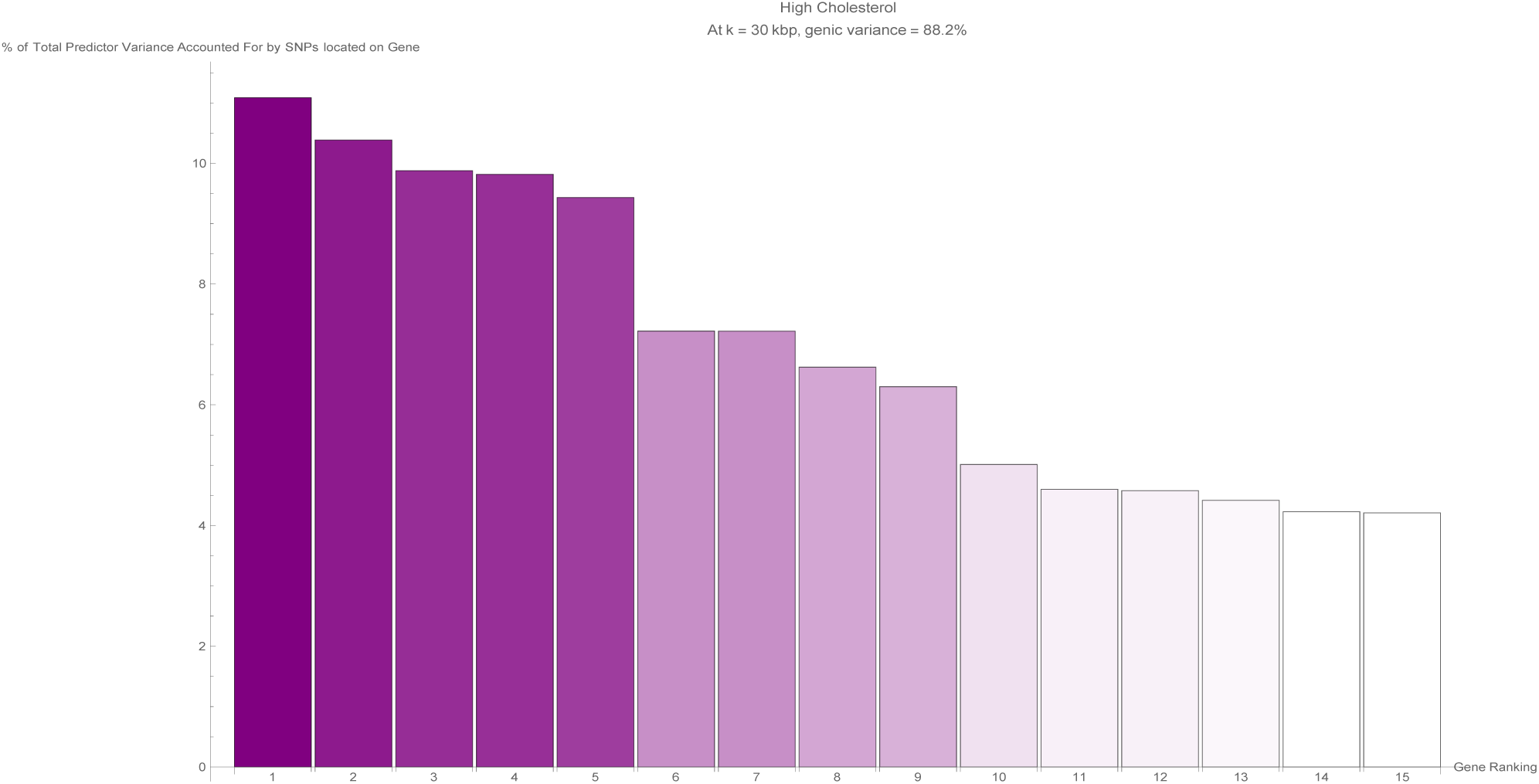
The fifteen largest total values of variance accounted for by predictor SNPs located on a single gene (in terms of the percentage of total variance accounted for by all predictor SNPs) for the high cholesterol predictor. Each vertical bar is colored violet with a depth of shade proportional to the height of the bar. Here, ‘genic’ SNPs are contained within the GENCODE Release 19 gene boundaries plus 30 kilo base pairs at both ends.

**Figure 23:**
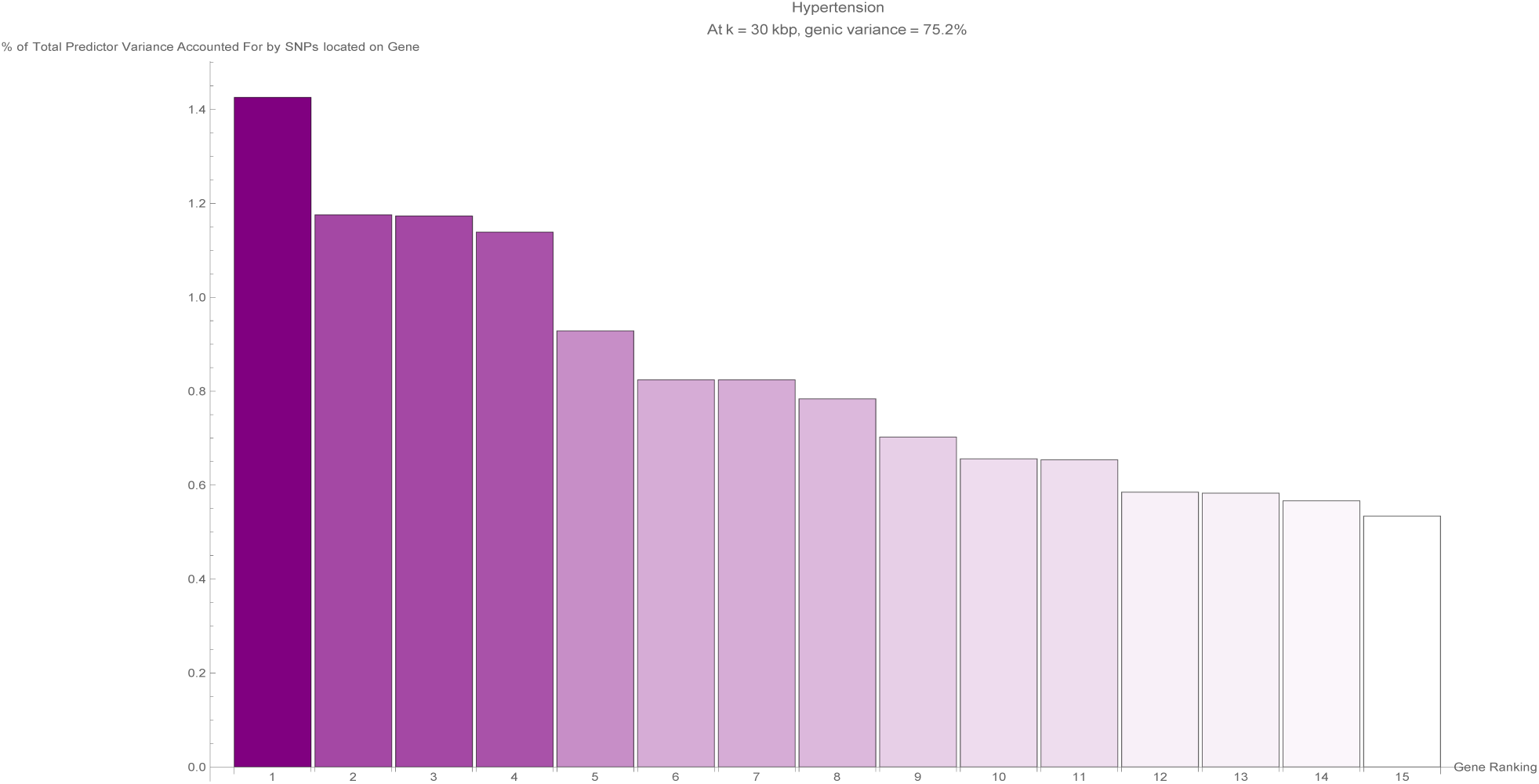
The fifteen largest total values of variance accounted for by predictor SNPs located on a single gene (in terms of the percentage of total variance accounted for by all predictor SNPs) for the hypertension predictor. Each vertical bar is colored violet with a depth of shade proportional to the height of the bar. Here, ‘genic’ SNPs are contained within the GENCODE Release 19 gene boundaries plus 30 kilo base pairs at both ends.

**Figure 24:**
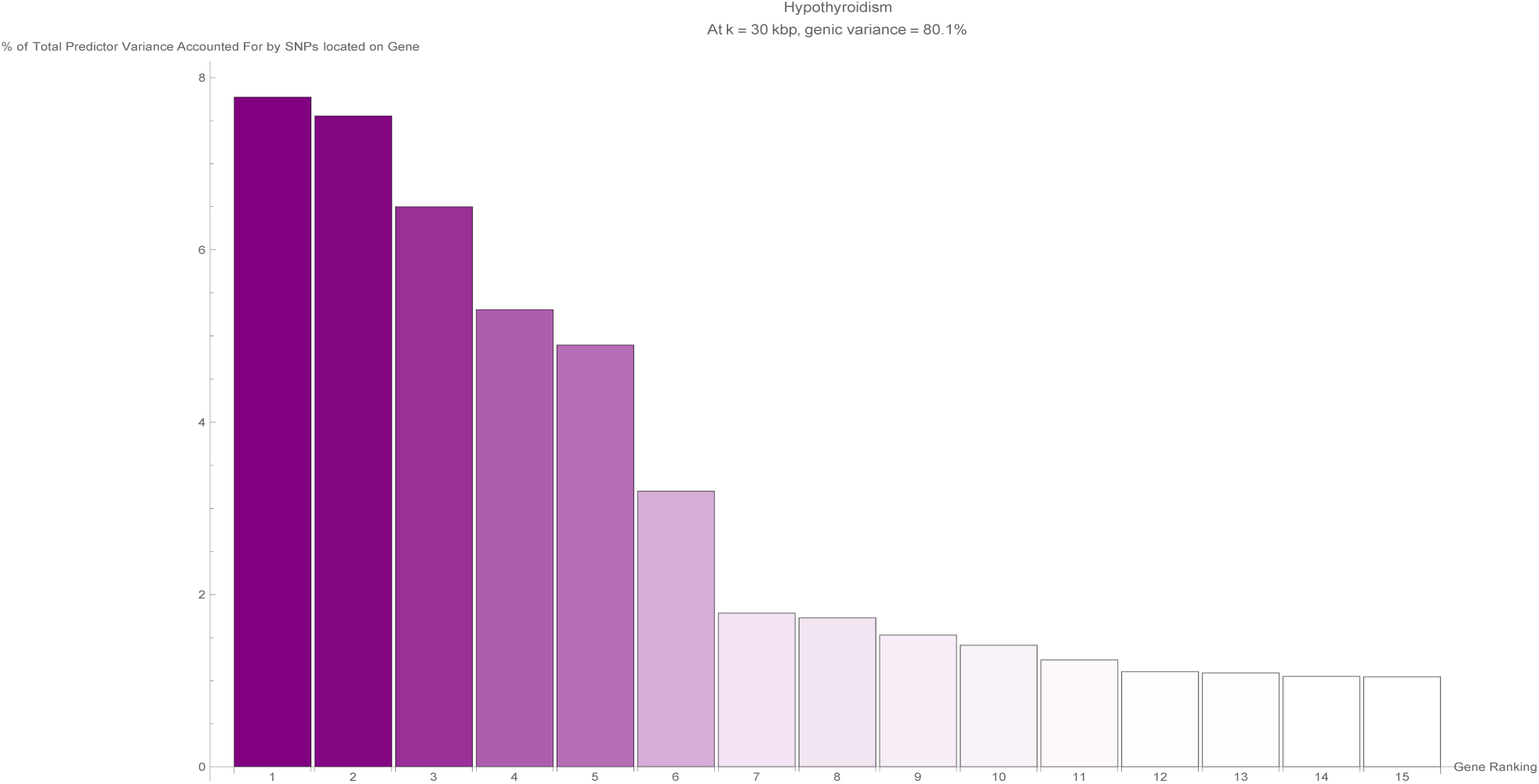
The fifteen largest total values of variance accounted for by predictor SNPs located on a single gene (in terms of the percentage of total variance accounted for by all predictor SNPs) for the hypothyroidism predictor. Each vertical bar is colored violet with a depth of shade proportional to the height of the bar. Here, ‘genic’ SNPs are contained within the GENCODE Release 19 gene boundaries plus 30 kilo base pairs at both ends.

**Figure 25:**
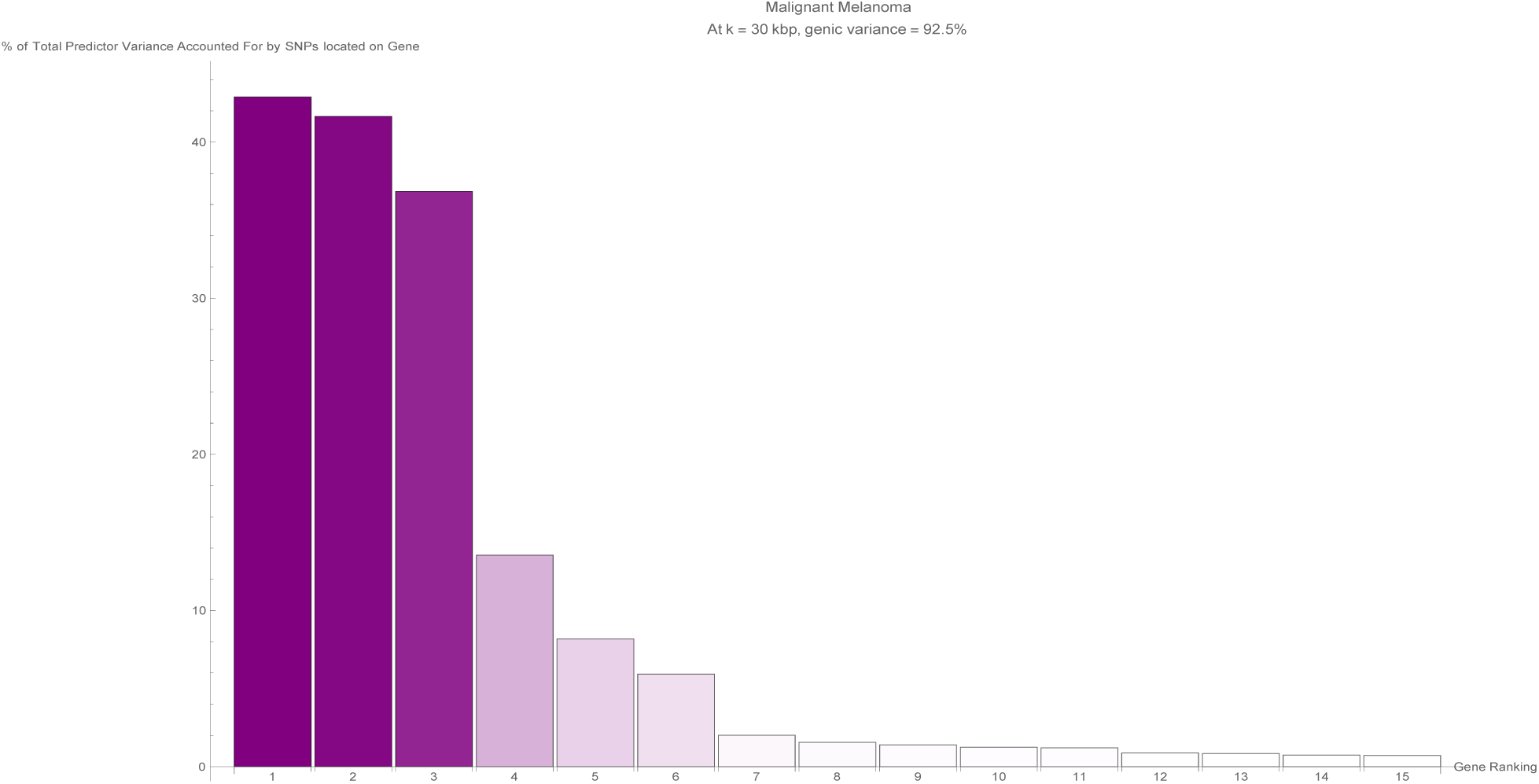
The fifteen largest total values of variance accounted for by predictor SNPs located on a single gene (in terms of the percentage of total variance accounted for by all predictor SNPs) for the malignant melanoma predictor. Each vertical bar is colored violet with a depth of shade proportional to the height of the bar. Here, ‘genic’ SNPs are contained within the GENCODE Release 19 gene boundaries plus 30 kilo base pairs at both ends.

**Figure 26:**
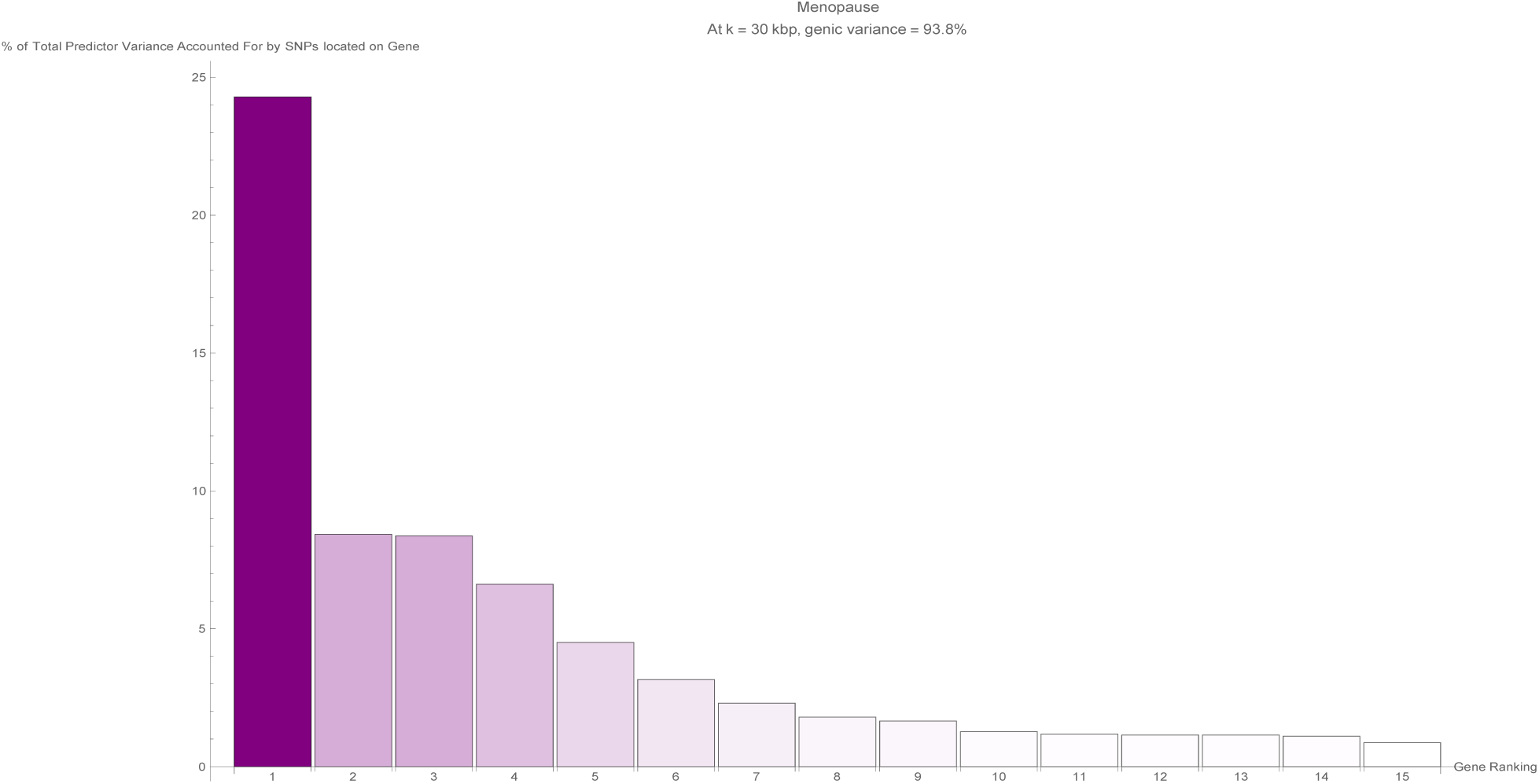
The fifteen largest total values of variance accounted for by predictor SNPs located on a single gene (in terms of the percentage of total variance accounted for by all predictor SNPs) for the menopause predictor. Each vertical bar is colored violet with a depth of shade proportional to the height of the bar. Here, ‘genic’ SNPs are contained within the GENCODE Release 19 gene boundaries plus 30 kilo base pairs at both ends.

**Figure 27:**
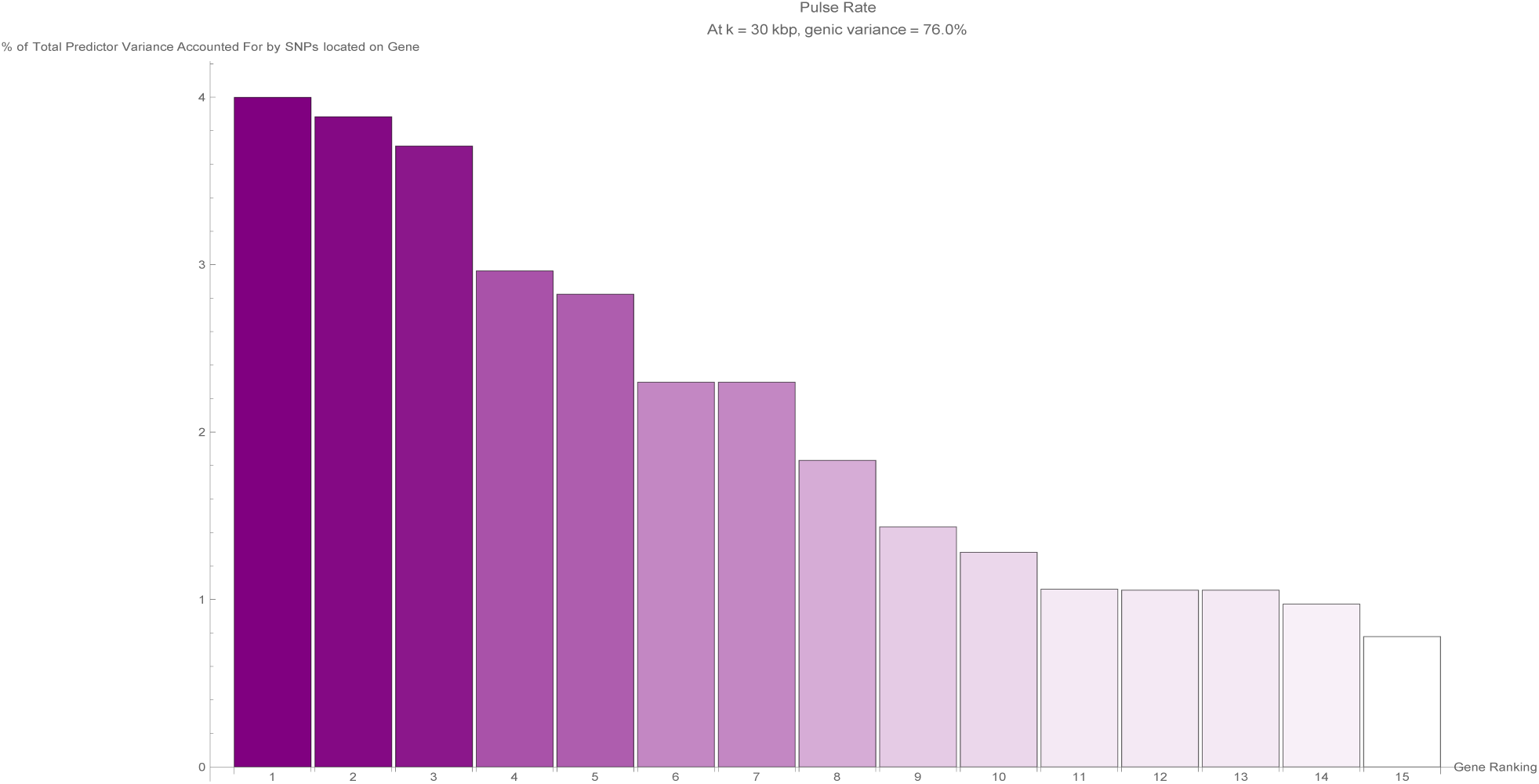
The fifteen largest total values of variance accounted for by predictor SNPs located on a single gene (in terms of the percentage of total variance accounted for by all predictor SNPs) for the pulse rate predictor. Each vertical bar is colored violet with a depth of shade proportional to the height of the bar. Here, ‘genic’ SNPs are contained within the GENCODE Release 19 gene boundaries plus 30 kilo base pairs at both ends.

**Figure 28:**
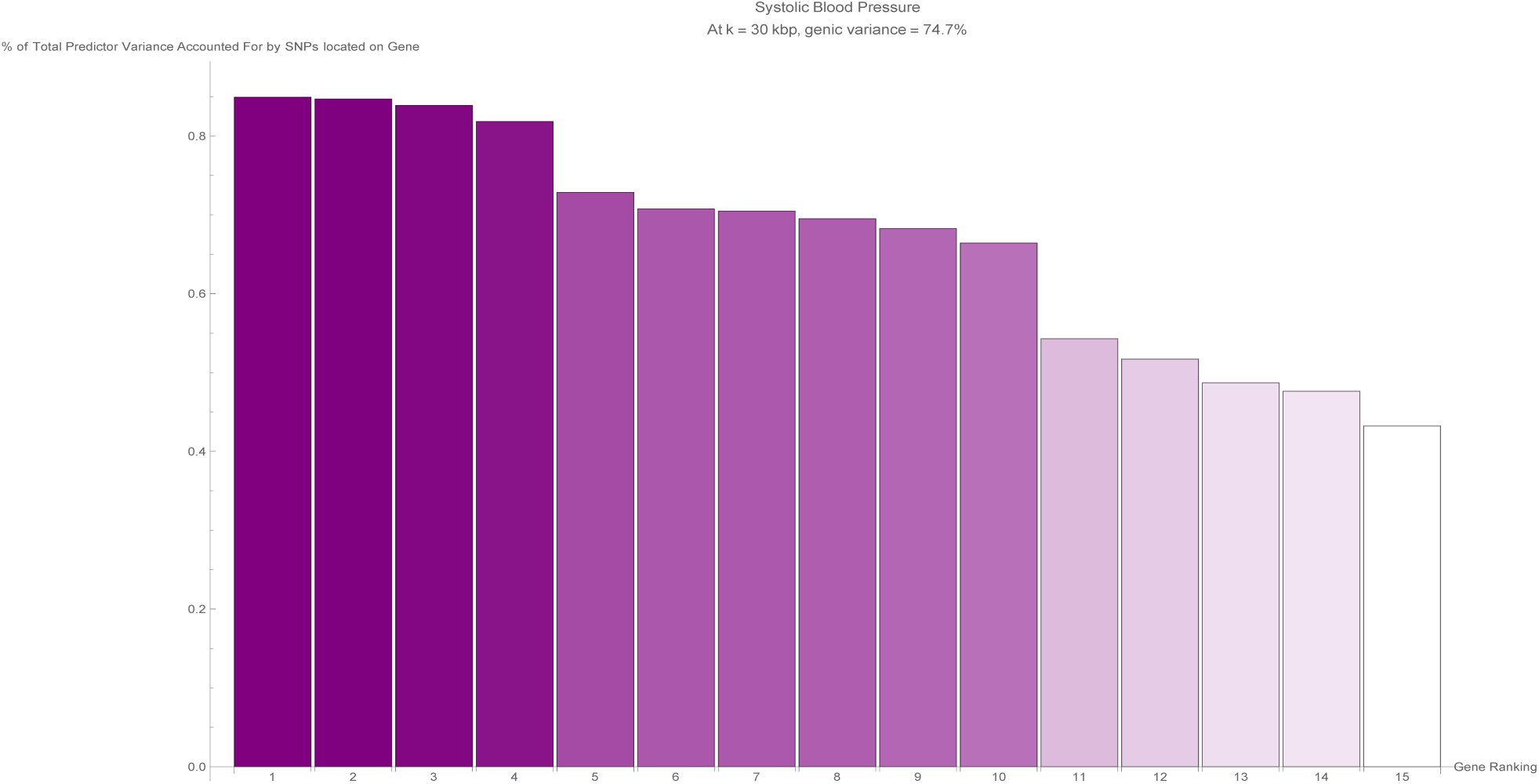
The fifteen largest total values of variance accounted for by predictor SNPs located on a single gene (in terms of the percentage of total variance accounted for by all predictor SNPs) for the systolic blood pressure predictor. Each vertical bar is colored violet with a depth of shade proportional to the height of the bar. Here, ‘genic’ SNPs are contained within the GENCODE Release 19 gene boundaries plus 30 kilo base pairs at both ends.

## Appendix B

Tables that list the top genes, as ordered by variance accounted for, for various phenotypes.

**Figure 29:**
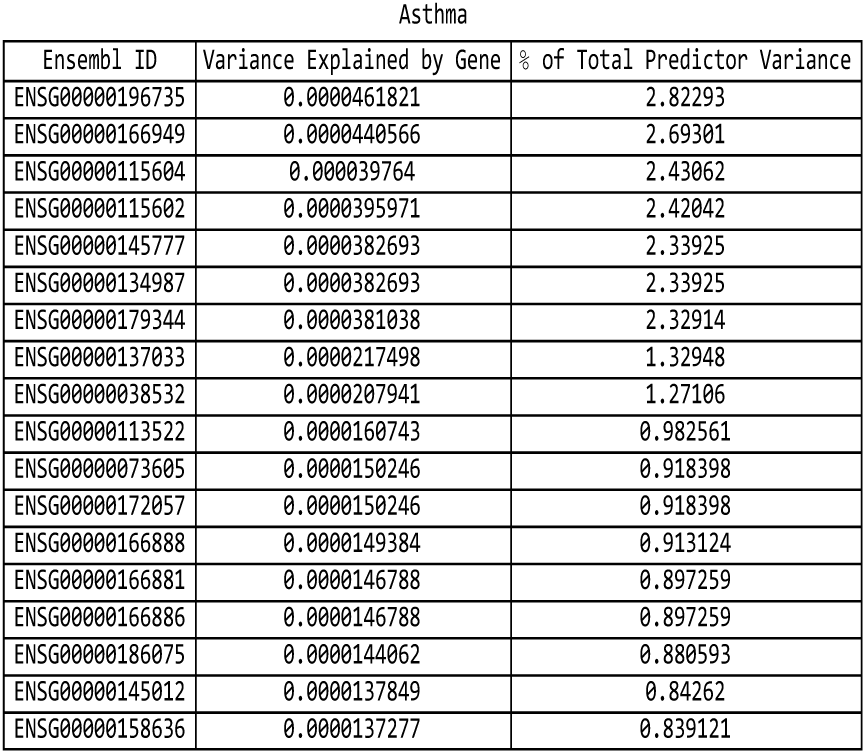
For the asthma predictor: list of genes responsible for the top fifteen values of variance accounted for by single genes, and the corresponding variance values, both explicit and expressed as a percentage of the total predictor variance.

**Figure 30:**
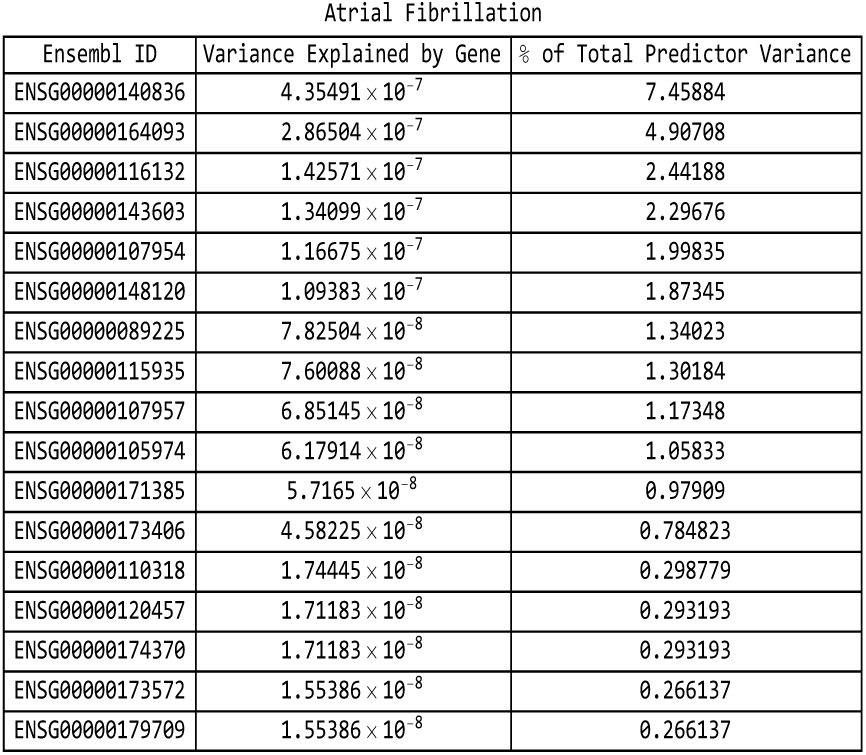
For the atrial fibrillation predictor: list of genes responsible for the top fifteen values of variance accounted for by single genes, and the corresponding variance values, both explicit and expressed as a percentage of the total predictor variance.

**Figure 31:**
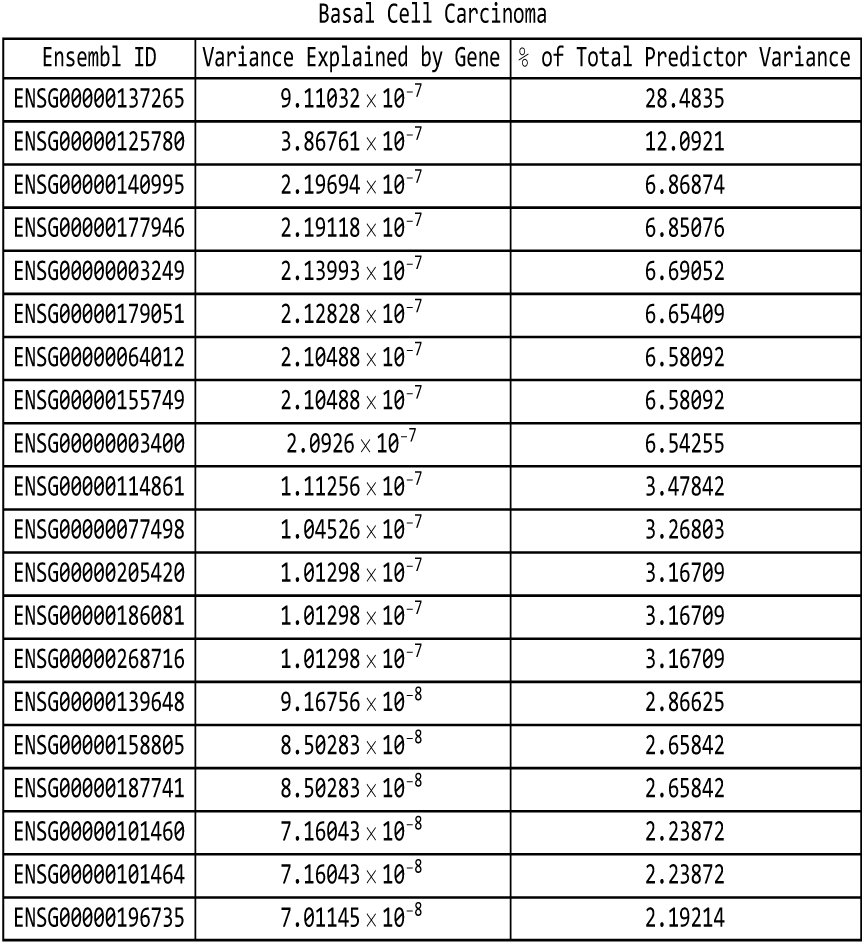
For the basal cell carcinoma predictor: list of genes responsible for the top fifteen values of variance accounted for by single genes, and the corresponding variance values, both explicit and expressed as a percentage of the total predictor variance.

**Figure 32:**
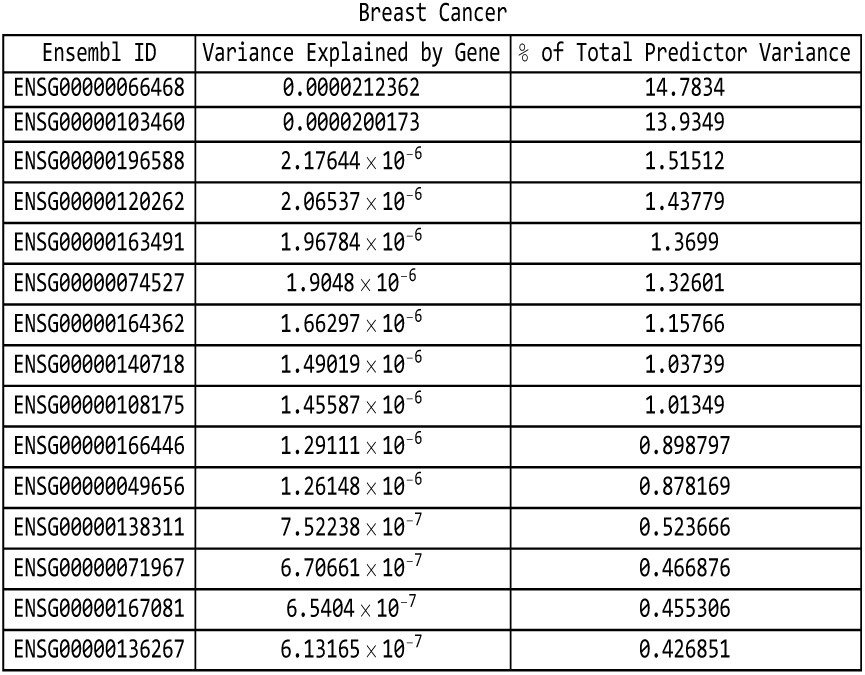
For the breast cancer predictor: list of genes responsible for the top fifteen values of variance accounted for by single genes, and the corresponding variance values, both explicit and expressed as a percentage of the total predictor variance.

**Figure 33:**
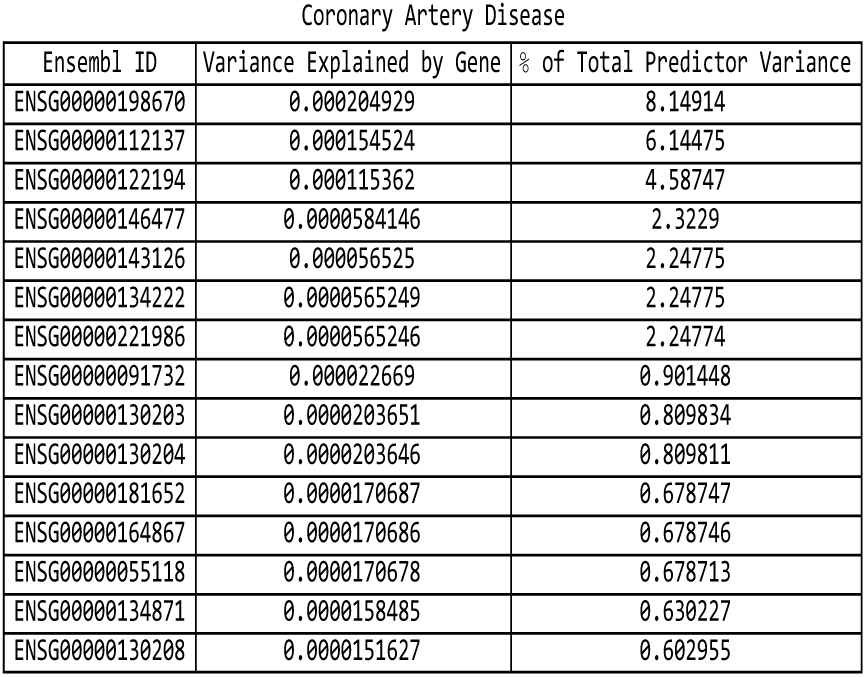
For the coronary artery disease predictor: list of genes responsible for the top fifteen values of variance accounted for by single genes, and the corresponding variance values, both explicit and expressed as a percentage of the total predictor variance.

**Figure 34:**
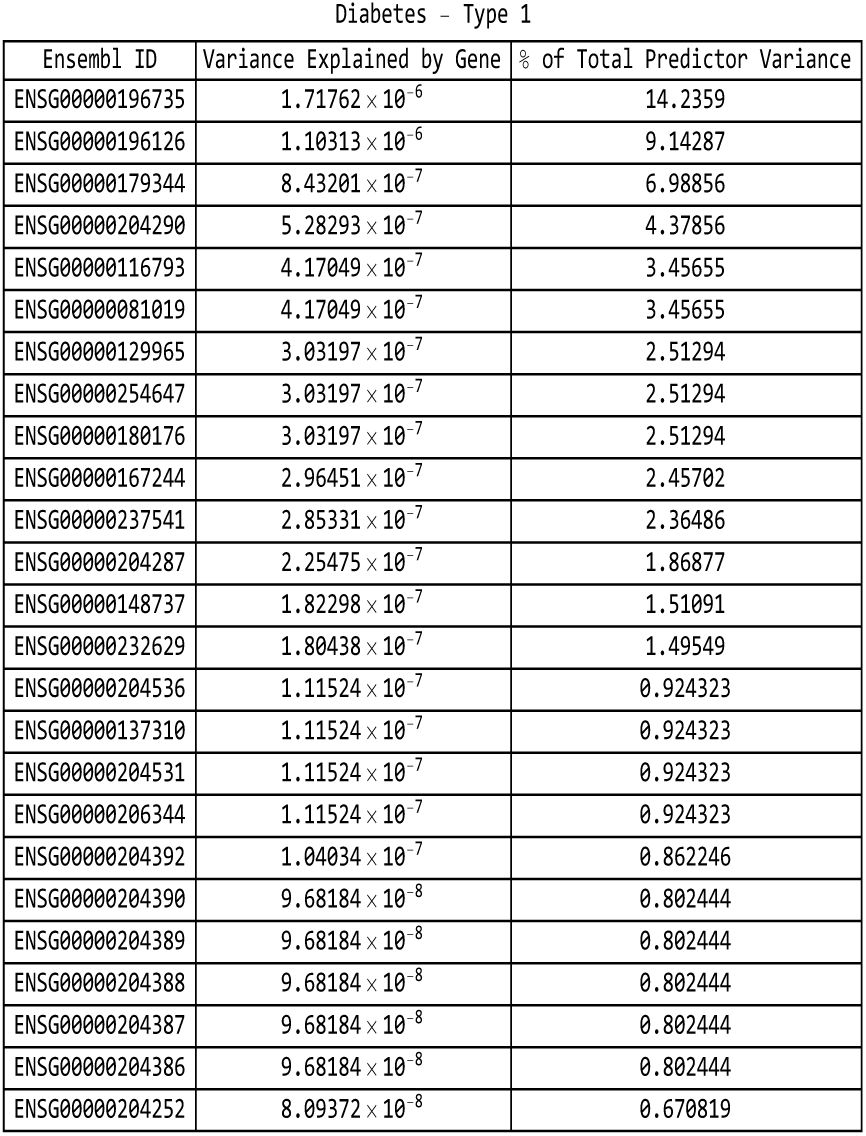
For the type-1 diabetes predictor: list of genes responsible for the top fifteen values of variance accounted for by single genes, and the corresponding variance values, both explicit and expressed as a percentage of the total predictor variance.

**Figure 35:**
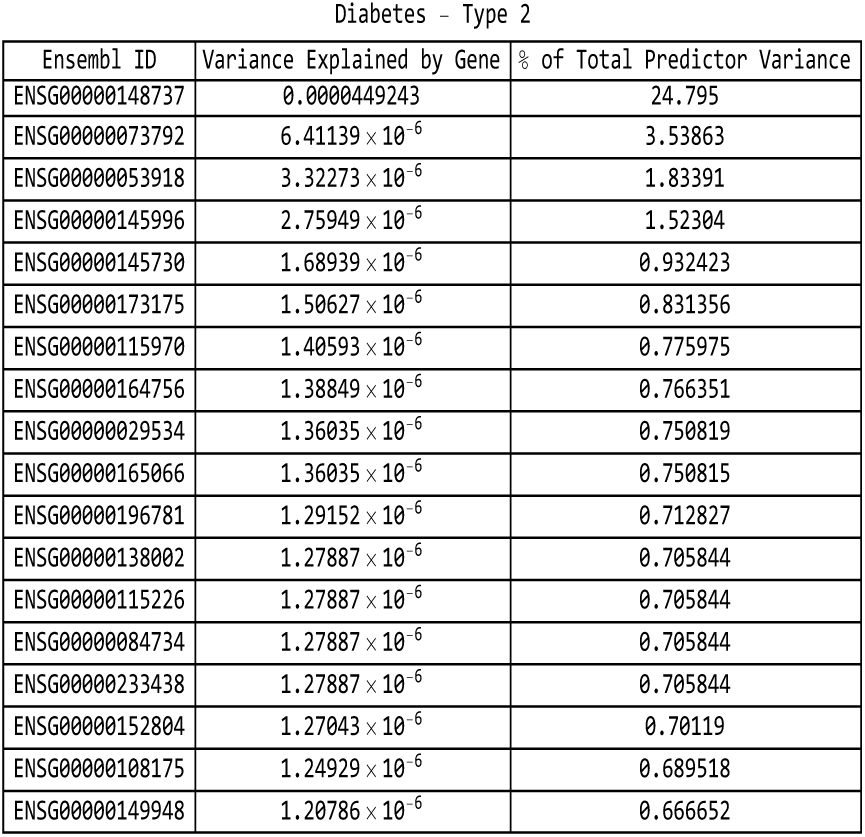
For the type-2 diabetes predictor: list of genes responsible for the top fifteen values of variance accounted for by single genes, and the corresponding variance values, both explicit and expressed as a percentage of the total predictor variance.

**Figure 36:**
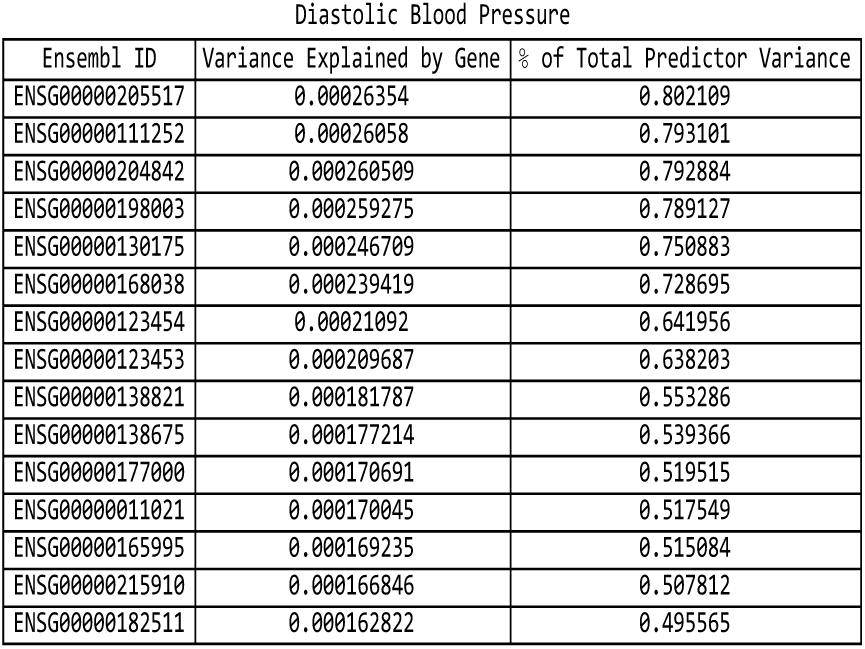
For the diastolic blood pressure predictor: list of genes responsible for the top fifteen values of variance accounted for by single genes, and the corresponding variance values, both explicit and expressed as a percentage of the total predictor variance.

**Figure 37:**
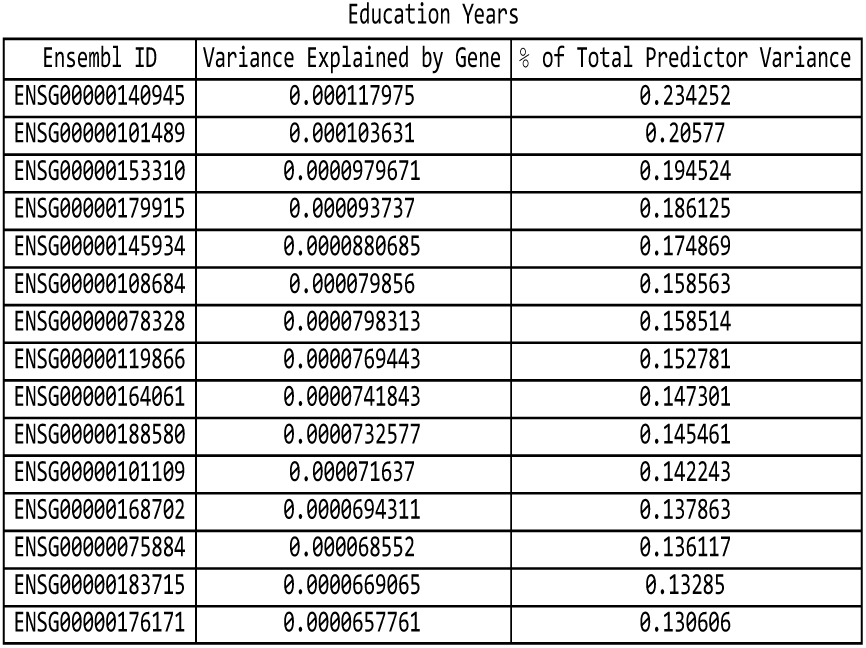
For the education years predictor: list of genes responsible for the top fifteen values of variance accounted for by single genes, and the corresponding variance values, both explicit and expressed as a percentage of the total predictor variance.

**Figure 38:**
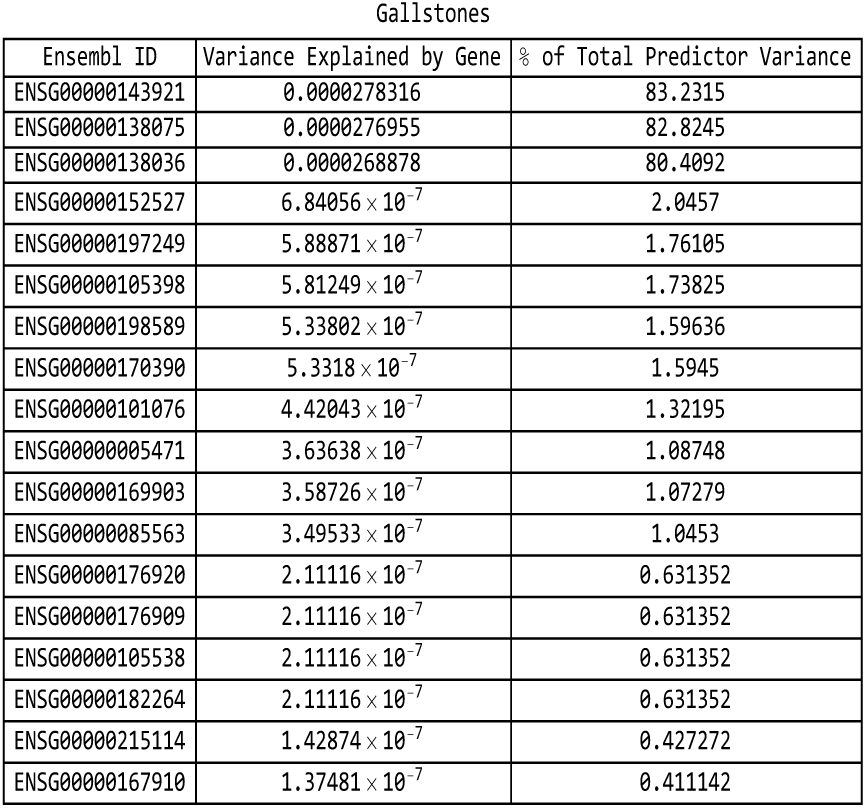
For the gallstones predictor: list of genes responsible for the top fifteen values of variance accounted for by single genes, and the corresponding variance values, both explicit and expressed as a percentage of the total predictor variance.

**Figure 39:**
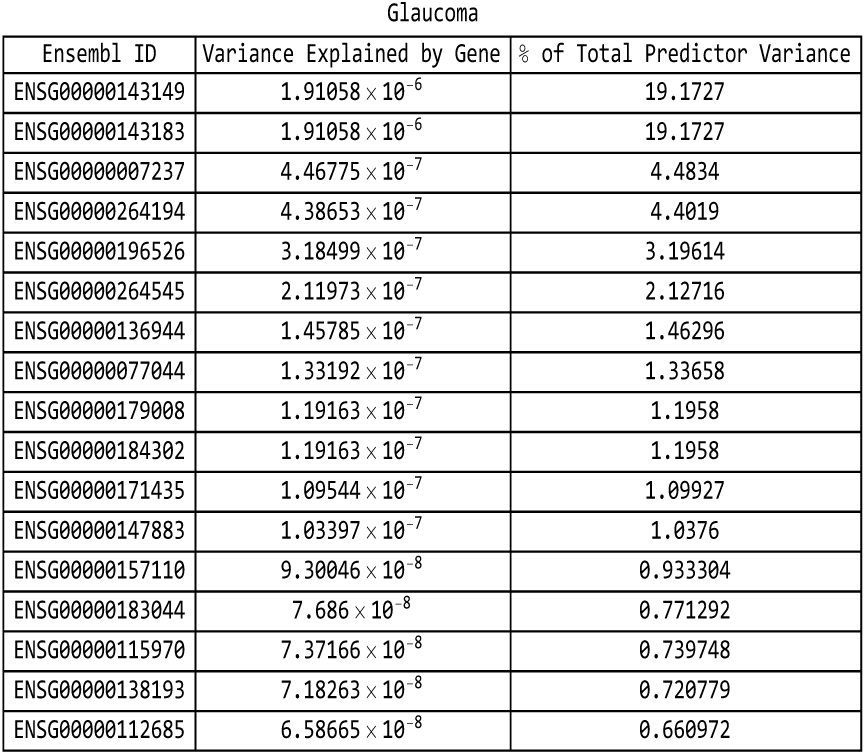
For the glaucoma predictor: list of genes responsible for the top fifteen values of variance accounted for by single genes, and the corresponding variance values, both explicit and expressed as a percentage of the total predictor variance.

**Figure 40:**
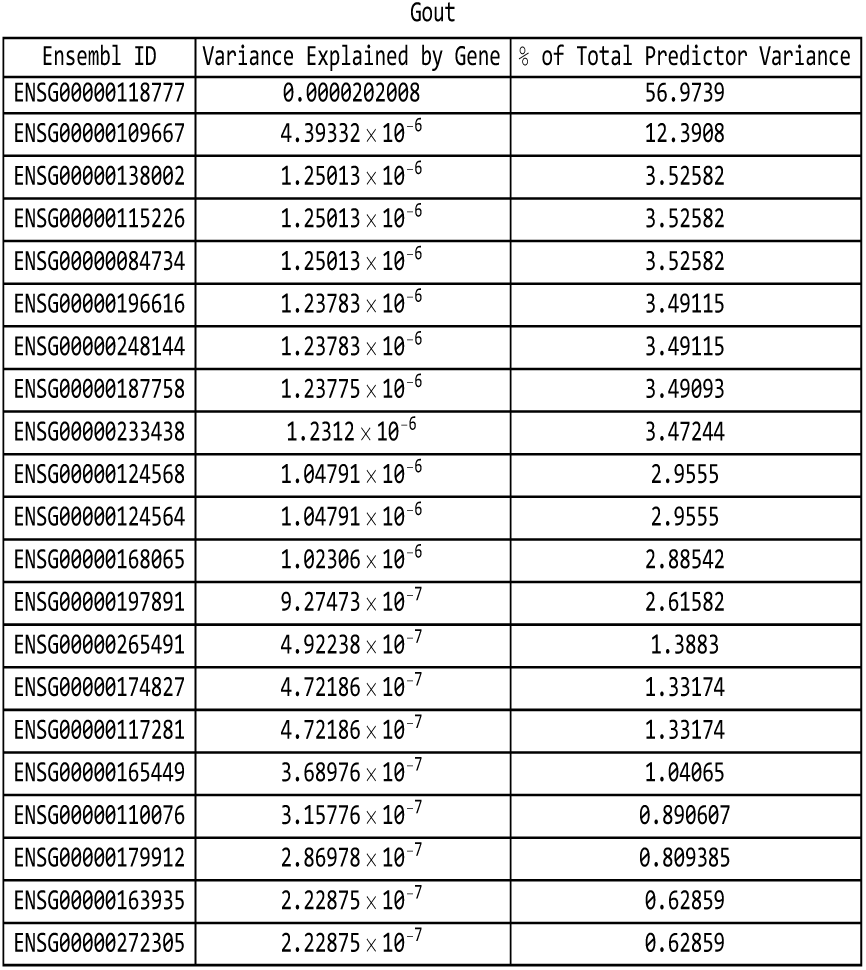
For the gout predictor: list of genes responsible for the top fifteen values of variance accounted for by single genes, and the corresponding variance values, both explicit and expressed as a percentage of the total predictor variance.

**Figure 41:**
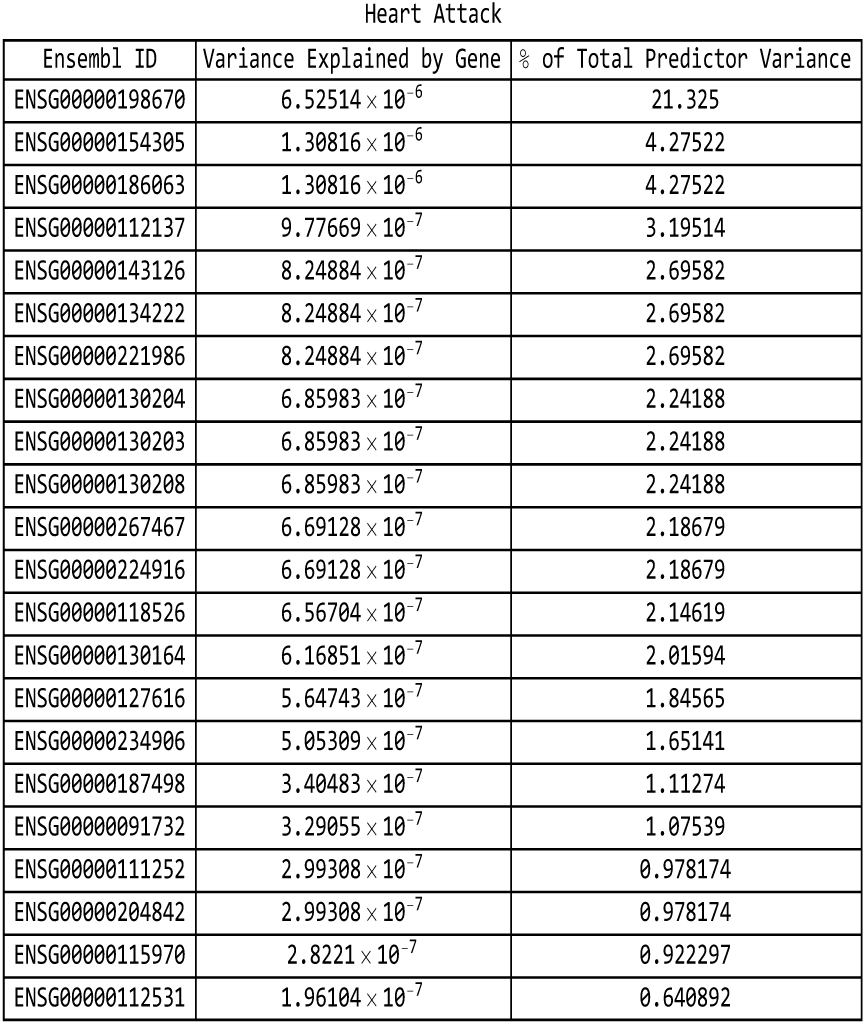
For the heart attack predictor: list of genes responsible for the top fifteen values of variance accounted for by single genes, and the corresponding variance values, both explicit and expressed as a percentage of the total predictor variance.

**Figure 42:**
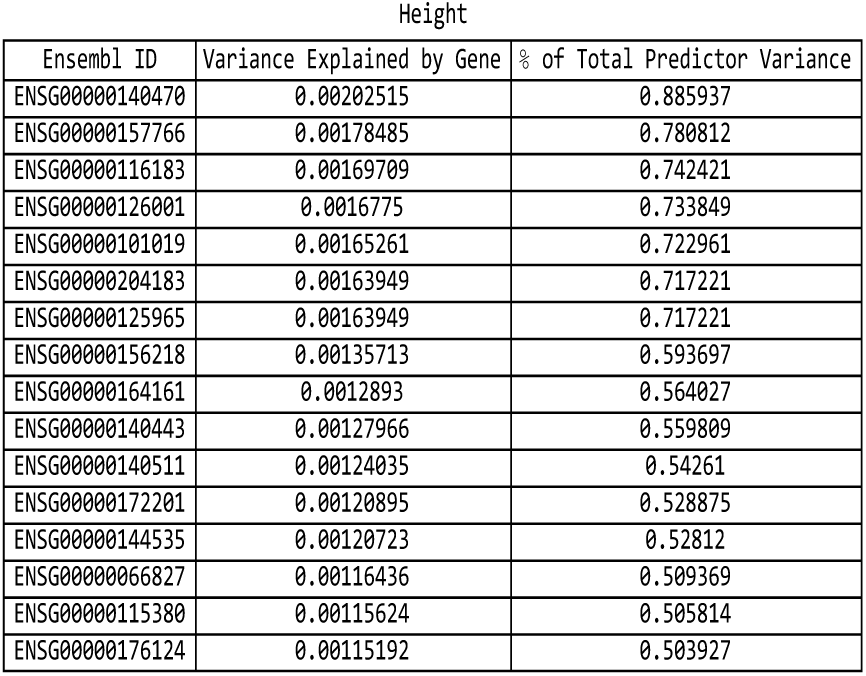
For the height predictor: list of genes responsible for the top fifteen values of variance accounted for by single genes, and the corresponding variance values, both explicit and expressed as a percentage of the total predictor variance.

**Figure 43:**
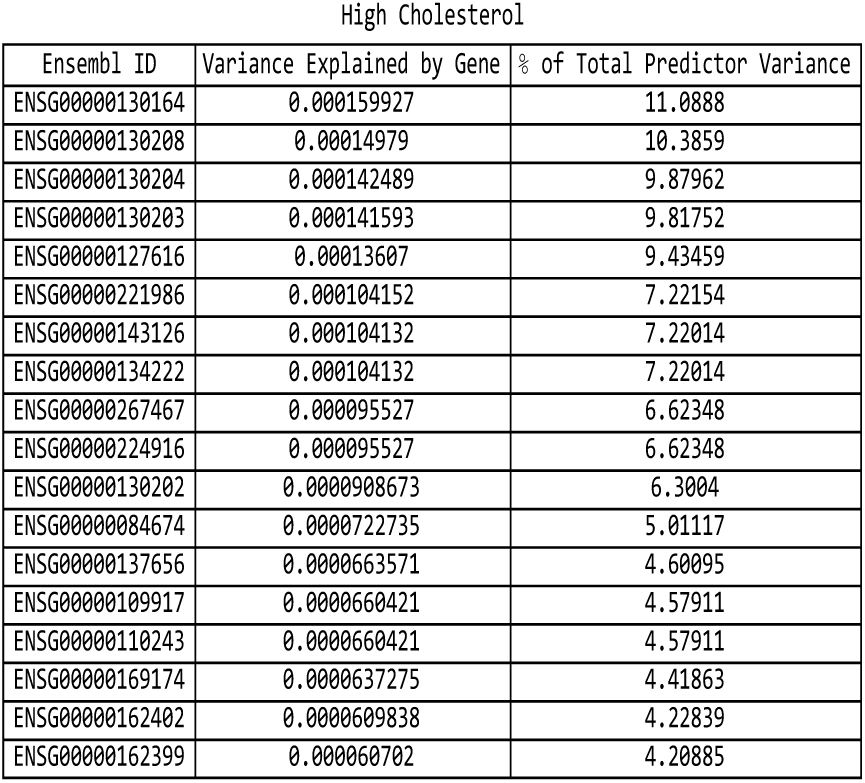
For the high cholesterol predictor: list of genes responsible for the top fifteen values of variance accounted for by single genes, and the corresponding variance values, both explicit and expressed as a percentage of the total predictor variance.

**Figure 44:**
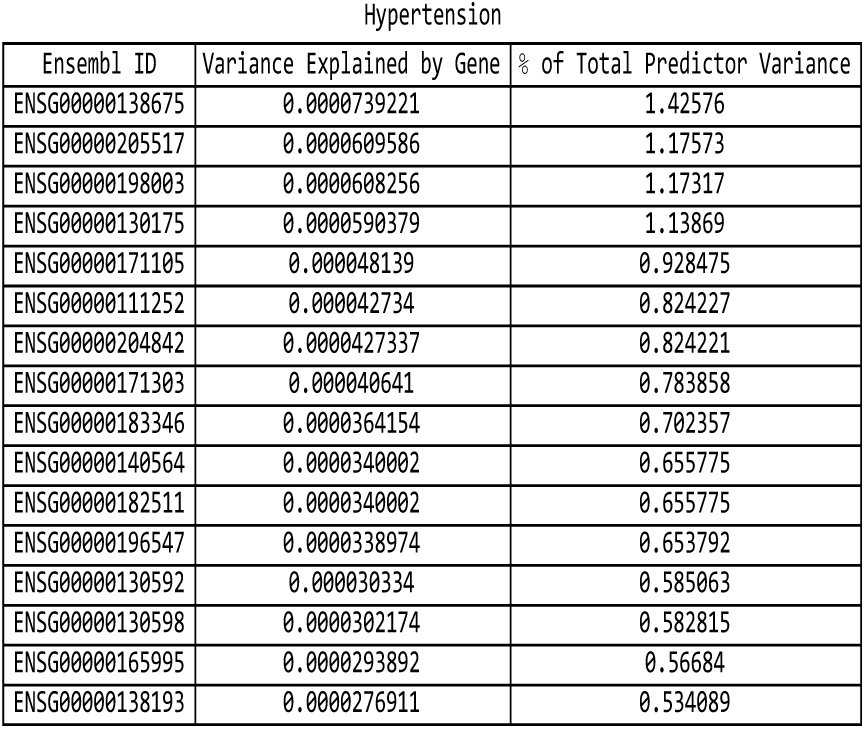
For the hypertension predictor: list of genes responsible for the top fifteen values of variance accounted for by single genes, and the corresponding variance values, both explicit and expressed as a percentage of the total predictor variance.

**Figure 45:**
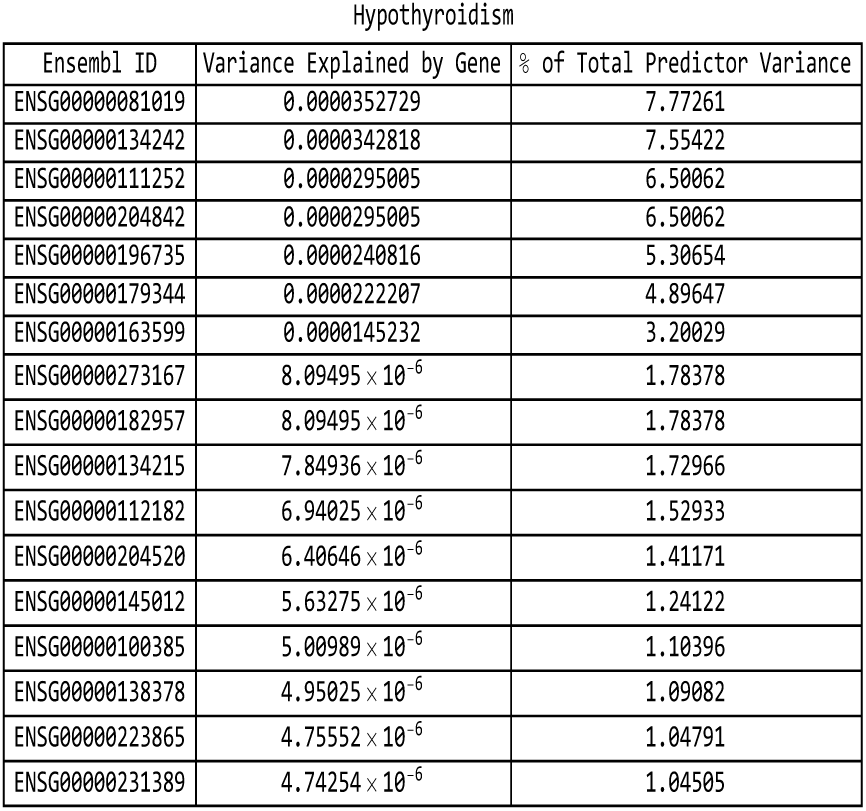
For the hypothyroidism predictor: list of genes responsible for the top fifteen values of variance accounted for by single genes, and the corresponding variance values, both explicit and expressed as a percentage of the total predictor variance.

**Figure 46:**
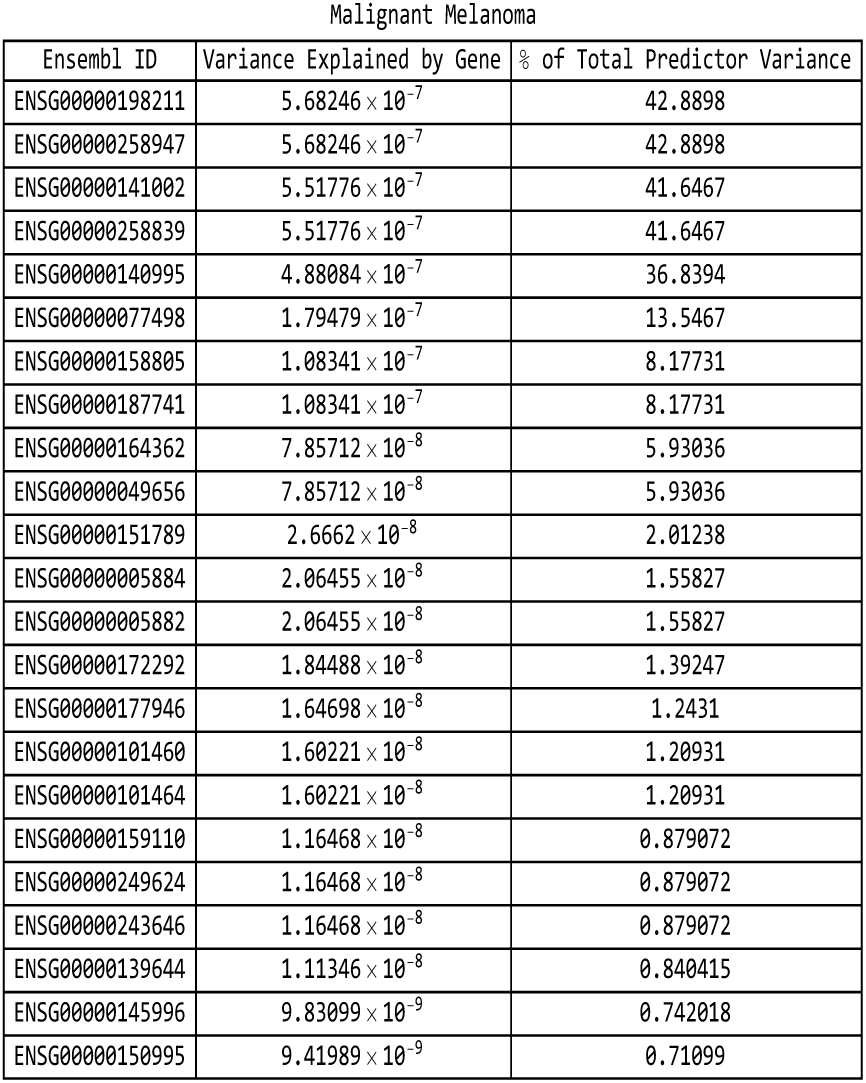
For the malignant melanoma predictor: list of genes responsible for the top fifteen values of variance accounted for by single genes, and the corresponding variance values, both explicit and expressed as a percentage of the total predictor variance.

**Figure 47:**
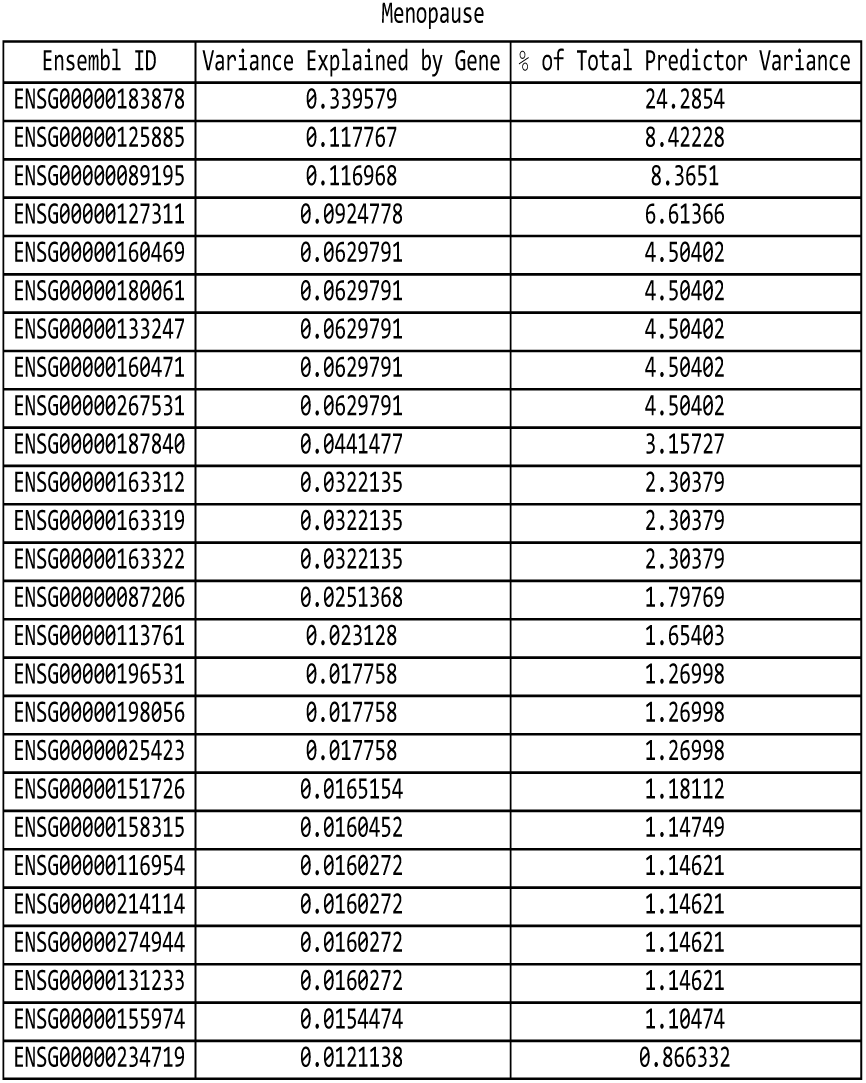
For the menopause predictor: list of genes responsible for the top fifteen values of variance accounted for by single genes, and the corresponding variance values, both explicit and expressed as a percentage of the total predictor variance.

**Figure 48:**
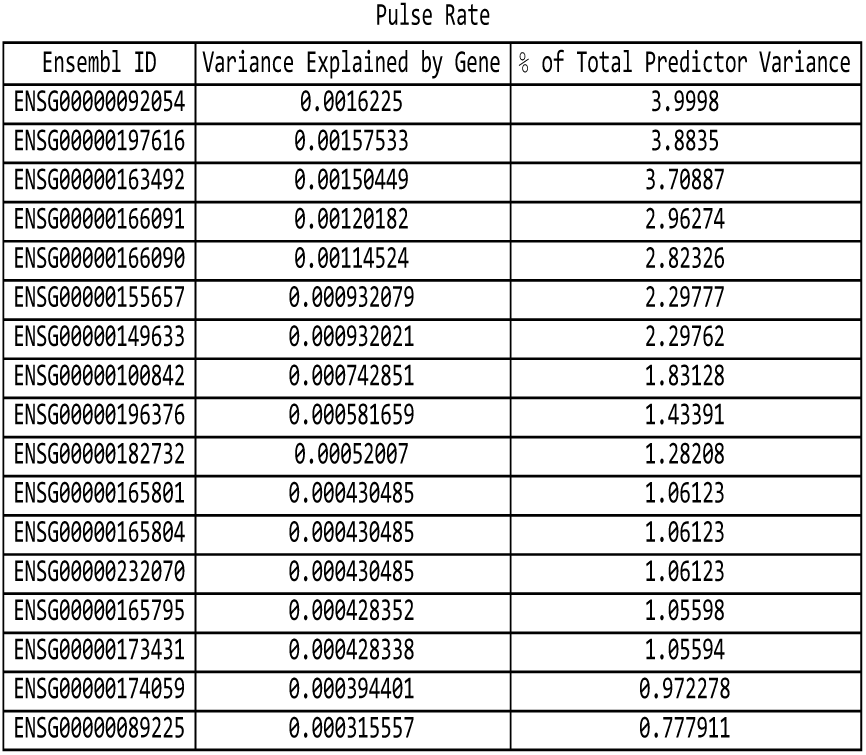
For the pulse rate predictor: list of genes responsible for the top fifteen values of variance accounted for by single genes, and the corresponding variance values, both explicit and expressed as a percentage of the total predictor variance.

**Figure 49:**
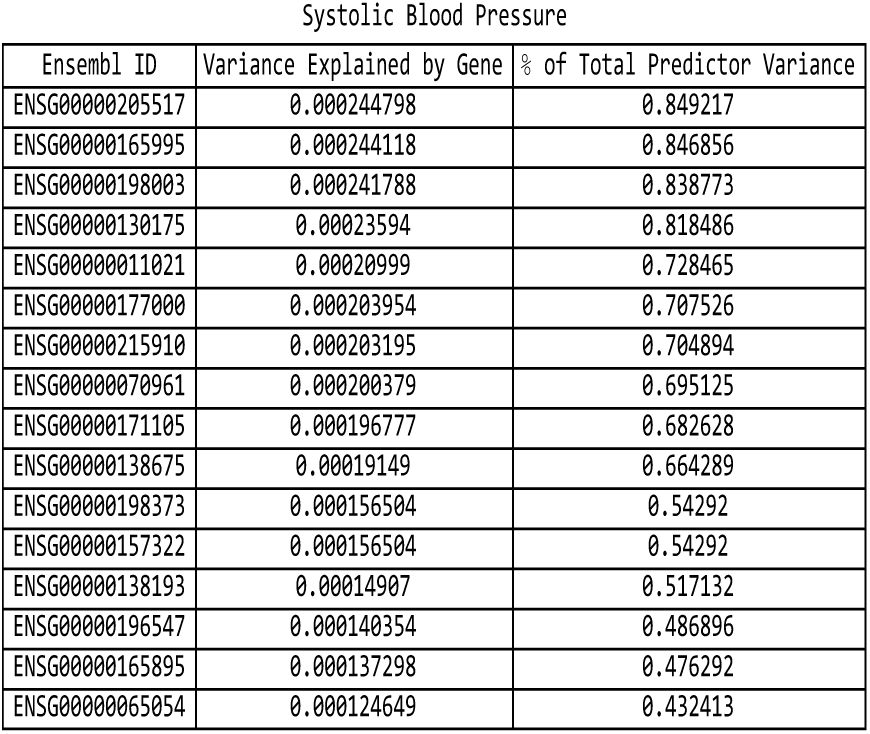
For the systolic blood pressure predictor: list of genes responsible for the top fifteen values of variance accounted for by single genes, and the corresponding variance values, both explicit and expressed as a percentage of the total predictor variance.

## Appendix C: UK Biobank Data

### C.1 Array Sequencing Data

Predictors (with the exception of the CAD predictor from [7]) were originally trained using the 2018 release of the UK Biobank. Predictor training was restricted to genetically British individuals (as defined using ancestry principal component analysis performed by UK Biobank) [35, 36]. In 2018, the UK Biobank re-released the dataset representing approximately 500,000 individuals genotyped on two Affymetrix platforms - approximately 50,000 samples on the UKB BiLEVE Axiom array and the remainder on the UKB Biobank Axiom array. The genotype information was collected for 488,377 individuals, and 805,426 SNPs, which were then subsequently imputed to a much larger number of SNPs. More details about the design of the array can be found on these documents from the UK Biobank website: an Axiom array content summary: http://www.ukbiobank.ac.uk/wp-content/uploads/2014/04/UK-Biobank-Axiom-Array-Content-Summary-2014-1.pdf, and the document detailing the Axiom array: http://www.ukbiobank.ac.uk/wp-content/uploads/2014/04/UK-Biobank-Axiom-Array-Datasheet-2014-1.pdf. Further information about the genotyping and phenotyping used to build the original predictors can be found in [5, 7].

### C.2 Exome Sequencing Data

In March 2019, the UK Biobank released whole-exome sequencing (WES) data for 49,960 participants [60]. Selection of participants for the study prioritized individuals with whole-body MRI imaging data from the UK Biobank Imaging Study, enhanced baseline measurements, hospital episode statistics (HES), linked primary care records, and admission to hospital with a primary diagnosis of asthma. In regards to age, sex and ancestry, the sequenced individuals are representative of the overall UK Biobank cohort. The sample set has 194 parent-offspring pairs, including 26 mother-father-child trios, 613 full-sibling pairs, 1 monozygotic twin pair and 195 second-degree genetically determined relationships.

Exomes were captured using a version of the IDT xGen Exome Research Panel v1.0. Multiplexed samples were sequenced with dual-indexed 75 × 75 bp paired-end reads on the Illumina NovaSeq 6000 platform using S2 flow cells. The specific genomic regions targeted for sequencing covered 39 megabases of the human genome, corresponding to 19,396 genes. In addition, the regions measuring 100 bp and located directly upstream and downstream of each target region were also sequenced.

A total of 4,735,722 variants located in targeted regions were identified. With adjacent (non-targeted) 100 bp regions included in the tally, a total of 9,693,526 indel and single nucleotide variants (SNVs) were observed. While only the target regions are required to meet all sequencing quality standards such as unique read coverage, variants in both target and adjacent regions were subjected to the same variant quality control metrics. Approximately 14% of coding variants identified via whole-exome sequencing were observed in the imputed sequence of 49,797 participants with both whole-exome sequencing and imputed data. 22.6% of the coding variants in the imputed data were not observed in the whole-exome sequencing data.

